# Visualization of stochastic expression of clustered protocadherin β isoforms *in vivo*

**DOI:** 10.1101/2025.03.04.641360

**Authors:** Ryosuke Kaneko, Yusuke Takatsuru, Manabu Abe, Yukiko U. Inoue, Ayako Morita, Satoko Horino, Takayoshi Inoue, Masahiko Watanabe, Kenji Sakimura, Yuchio Yanagawa, Takeshi Yagi

**Affiliations:** KOKORO-Biology Group, Laboratories for Integrated Biology, Graduate School of Frontier Biosciences, Osaka University, 1-3 Yamadaoka, Suita, Osaka 565-0871, Japan; Bioresource center, Gunma University Graduate School of Medicine, Maebashi, 371-8511, Japan; Medical Genetics Research Center, Nara Medical University, Kashihara, Nara 634-8521, Japan; Department of Nutrition and Health Science, Toyo University, Gunma 374-0193, Japan; Department of Cellular Neurobiology, Brain Research Institute, Niigata University, Niigata, Niigata 951-8585, Japan; Department of Biochemistry and Cellular Biology, National Institute of Neuroscience, National Center of Neurology and Psychiatry, Ogawahigashi, 4-1-1, Kodaira, Tokyo 187-8502, Japan; Department of Anatomy, Graduate School of Medicine, Hokkaido University, Sapporo, Hokkaido 060-8638, Japan; Department of Genetic and Behavioral Neuroscience, Gunma University Graduate School of Medicine, Maebashi, Gunma 371-8511, Japan

## Abstract

The stochastic expression of clustered protocadherin (cPcdh) establishes a single-cell identity fundamental to cellular self/non-self-discrimination. However, it has been challenging to reveal the spatiotemporal patterning of the stochastic cPcdh expression *in vivo*. We developed XFP (tdTomato or GFP) knock-in mice using a new strategy to enhance XFP expression, which allows us to visualize cPcdhβ3 or 19-positive cells throughout the brain. These mouse lines demonstrate the cell-type selectivity, spatial biases, inter-individual differences, left-right asymmetry, developmental regulation, alteration in pathological or aging brain, and monoallelic expression of stochastic cPcdhβ expression *in vivo*. Our findings further demonstrate that the cPcdhβ3 expression undergoes significant changes in the mature brain over time. These results demonstrate the potential of these reporter mice to advance our understanding of cPcdh cellular barcode *in vivo*.

**One Sentence Summary:** Novel established cPcdhβ reporter mouse lines reveal stochastic expression of cPcdhβ isoforms *in vivo*.

## Main Text

Gene expression programs with precise spatial and temporal regulations provide a foundation for the development of multicellular organisms. While most genes are deterministically regulated, there are systems in the animal kingdom in which stochastic gene expression is preferred. This is because it provides diverse cellular identities that are challenging to achieve through deterministic gene regulation(*1*, *2*). For instance, the production of immunoglobulins by VDJ recombination generates near-complete random diversities in the immune system. Stochastic gene expression is also responsible for generating diverse cellular identities in the nervous system. This impact on higher brain function is evident in two notable examples, namely mammalian olfactory receptors and the *Drosophila* Down syndrome cell adhesion molecule (Dscam1) gene. The stochastic expression of olfactory receptors ensures proper olfactory information processing according to spatial patterns in the mouse olfactory epithelium. The stochastic expression of Dscam1 is essential for the normal patterning of axons and dendrites in the fly brain.

The stochastic expression of clustered protocadherin (cPcdh) family gene provides diverse cellular identities in vertebrate central nervous systems. The mammalian cPcdh family gene, first identified by Kohmura et al. in 1998 as CNRs(*3–6*), consists of three gene clusters (cPcdha, cPcdhb, and cPcdhg) located in tandem on a single chromosome. There are 14, 22, and 22 isoforms in the mouse, respectively(*7*, *8*). Each cPcdh cluster is independently regulated, and the cPcdh repertoires in each neuron are generated by stochastic gene regulation, comprising random promoter choice and DNA methylation(*9–12*). This stochastic cPcdh expression confers a highly diverse cellular barcode to individual neurons, which then form cis-homodimers or cis-heterodimers that participate in strict homophilic interactions at the cell surface. These interactions are essential for cellular self/non-self discrimination(*12–17*).

The cPcdh cellular barcode plays an important role in the mammalian brain, but its physiological roles have been proven for only a few cell types so far. The main reason for this is the technical difficulties involved in revealing the temporal and spatial patterning of this identity *in vivo*(*13–15*, *18–20*). For example, we and others have shown the spatial patterning of cells expressing a given cPcdh isoform using *in situ* hybridization or immunostaining experiments(*15*, *21–25*). However, because of those higher homologies, the possibility of cross-detecting other cPcdh mRNAs/proteins cannot be excluded. Furthermore, despite repeated attempts, visualization of stochastic cPcdh expression has not been successful in cPcdh reporter mic. Moreover, available techniques can only be applicable to fixed or dissociated tissues, therefore there is no method that allows the direct visualization of endogenous cPcdh-expressing cells in the living brain.

Here, we developed a more specific approach to determine the spatiotemporal distribution of cPcdh-positive cells in fixed and living brains. By modifying endogenous cPcdhβ clusters, we generated several knock-in mouse lines expressing fluorescent protein cDNAs (XFP; tdTomato or GFP) under the control of endogenous cPcdhβ3 or cPcdhβ19 expressional regulation. In these reporter mouse lines, cells that translate the modified cPcdhβ3 or cPcdhβ19 alleles can be detected by fluorescent proteins. We revealed that cells expressing a given cPcdhβ isoform are stochastically but spatially biased with cell-type dependency throughout the brain and is influenced by brain dysfunction. Unexpectedly, cPcdhβ expression in neurons/glia changes within weeks in the living brain. The cPcdhβ reporter mice provide unique models to investigate the spatiotemporal distributions and physiological roles of cPcdh cellular barcode *in vivo*.

## Results

### Modification of the endogenous cPcdhβ3 to visualize its expression *in vivo*

We aimed to develop a genetic approach for visualizing stochastic cPcdh expression *in vivo*. We firstly choose the cPcdhβ3 loci as a model because (1) cPcdhβ is technically easier than cPcdhα and cPcdhγ genes as the cPcdhβs are single-exon genes rather than spliced genes, and (2) it has been proven to have stochastic expression patterns by both *in situ* hybridization and single-cell RT-PCR experiments (Fig. S1)(*8*, *15*).

To ensure that cells expressing fluorescent protein simultaneously expressed a cPcdhβ protein, we designed a knock-in locus that expressed both cPcdhβ3 and tdTomto. To enhance fluorescence intensity, we developed a multiple tandem 2A peptide-flanked tdTomato (multi-tandem tdTomato), and it showed the enhanced fluorescence intensity upon transient expression in Cos7 cells (Fig. S2A). Thus, we knocked-in the multi-tandem tdTomato into the endogenous β3 locus (Fig. 1A-B and Fig. S2B-C). Both the heterozygous and homozygous knock-in mice were fertile and did not exhibit any morphological or developmental defects.

**Fig. 1.**
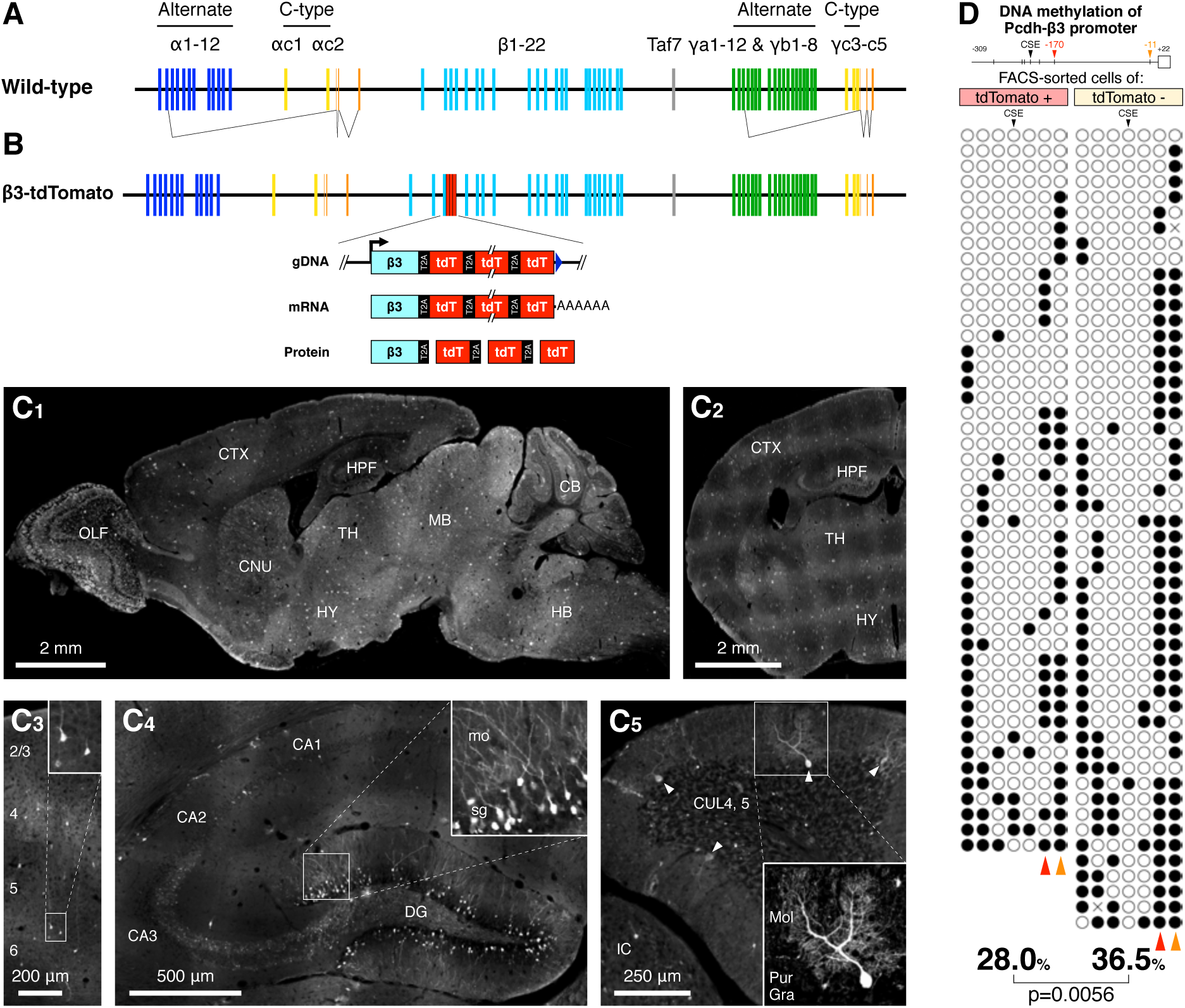
Generation, visualization, and validation of the cPcdhβ3-tdTomato reporter mouse. (A)Schematic representation of the mouse clustered cPcdh genomic locus. The cPcdha alternate, cPcdhb, cPcdhg alternate genes (Blue, Aqua, and Green, respectively) show stochastic expression in several neuron types, whereas c-type genes from cPcdh and clusters (yellow) are constitutively expressed in Purkinje cells. Orange: constant region exons of cPcdha and cPcdhg clusters. Gray: Taf7, non-Pcdh genes. (B) Genomic structure of cPcdhβ3-tdTomato reporter alleles, which includes multi-tandem tdTomato in the endogenous cPcdhβ3 locus. (C) tdTomato expression from the cPcdhβ3-tdTomato reporter alleles (4-week-old) in the brain sagittal section (C1), the forebrain coronal section (C2), the cerebral cortex coronal section (C3), the hippocampus coronal section (C4), and the cerebellum sagittal section. The boxed region is magnified in C3, C4, and C5. (D) Validation of Pcdhβ3-tdTomato reporter. DNA methylation analysis of the cPcdhβ3 promoter in the tdTomato (+) and (-) cells purified from E16.5 neocortex of the cPcdhβ3-tdTomato reporter allele. Red and orange arrows represent CpG positions showing significantly lower methylation in tdTomato (+) cells.

The endogenous cPcdhβ3 is stochastically expressed in adult brains (Fig. S1). Therefore, we examined whether the β3-tdTomato reporter mice recapitulated the endogenous expression of cPcdhβ3 by histological examination. As expected, stochastic expression of tdTomato was confirmed 4-weeks-old brain (Fig. 1C). Stochastic tdTomato expression was observed in previously unknown regions, such as the caudate putamen, thalamus, superior colliculus, inferior colliculus, and brainstem. The frequency of tdTomato^+^ Purkinje cells is comparable to our previous data: 2.4 ± 0.2% for the reporter and 3.6% for endogenous cPcdhβ3 obtained by single-cell RT-PCR(*15*). The overall distribution of cells expressing tdTomato seems random, consistent with previous results, validating our reporter mouse line(*15*).

A reporter mouse expressing a fluorescent protein offers the advantage of allowing the purification of fluorescence positive cells using a fluorescence-activated cell sorter (FACS). We next asked whether β3-tdTomato reporter mice could enrich cPcdhβ3-expressing cells using FACS. The cerebral cortex of E16.5 embryonic mice were micro-dissected and subjected to enzymatic digestion to obtain a single-cell suspension, followed by cell sorting. To verify that the purified tdTomato-positive cells are cPcdhβ3-expressing cells, we analyzed their DNA methylation of cPcdhβ3 promoter by bisulfite sequencing, since cPcdh expression is inverse correlated with promoter DNA methylation(*9*, *27*). The DNA methylation of cPcdhβ3 promoter in the tdTomato^+^ was significantly lower than that of tdTomato^-^ cells, while the DNA methylation of cPcdhβ8 and cPcdhβ19 promoter were comparable between the tdTomato^+^ and tdTomato^-^ cells (Fig. 1D and Fig. S3). The results confirm that the purified tdTomato^+^ population is highly enriched with cPcdhβ3-expressing cells. Taken together, β3-tdTomato reporter mice can be used reliably to characterize the cPcdhβ3-expressing cells *in vivo*.

### β3-tdTomato reporter mouse reveals cell-types showing stochastic cPcdhβ3 expression

Although stochastic cPcdhα, cPcdhβ, and cPcdhγ expression is fundamental for establishing the cPcdh single-cell identity, limited information regarding the cell types that stochastically express cPcdh has hampered research progress in understanding its roles in the brain. Here, we identified the cell-types showing stochastic cPcdhβ3 expression in the brain by double immunofluorescence of tdTomato and cell-type markers (Fig. S4). As shown by previous studies, developing hippocampus, retina, olfactory epithelium, and olfactory bulb demonstrated tdTomato expression in a subset of neurons, further validating our reporter mouse line (Fig. S4A-D). The markers included GABA for inhibitory interneurons, parvalbumin (PV) and somatostatin (SST) for interneuron subtypes, Pax6 for neural stem/progenitor cells in the ventricular zone, Tryptophan Hydroxylase 2 (TPH2) for serotonergic neuron, tyrosine hydroxylase (TH) for dopaminergic and noradrenergic neurons, GFAP for astrocyte, GST-π for oligodendrocyte, and IbaI for microglia. The photographs in Fig. S4E-M clearly show that a subset of all cell types, including neural stem/progenitor cells, astrocytes, and oligodendrocytes, expressed tdTomato, suggesting the presence of diverse cPcdh single-cell identity in these cell types. However, the photographs showed no tdTomato expression in microglia or blood vessels, suggesting that cPcdhβ is not expressed in these cells (Fig. S4N).

We also examined the cPcdhβ expression in non-neural organs (Fig. S4O and S4P). Kidney section from the β3-tdTomato reporter mouse showed no tdTomato expression, whereas the testis section showed rather constitutive expression of tdTomato in differentiated sperm(*28*), clearly demonstrating the cPcdhβ expression is not always stochastic. These data revealed that the cPcdhβ expression pattern depends on cell-types and organ, showing the usefulness of examining the cPcdhβ expression at single-cell resolution using the β3-tdTomato reporter mouse.

### cPcdhβ3 expression shows spatial-bias, inter-individual variation and left-right asymmetry

We next examined the spatial distribution of cPcdhβ3 expression using cerebellar Purkinje cells as a model (Fig. 2). First, we investigated whether cPcdhβ3 expression was spatially biased by examining the regional differences in stochastic cPcdhβ3 expression in cerebellar Purkinje cells (Fig. 2A-D). Since the Purkinje cells are subdivided into two main groups based on Zebrin II (also known as Aldoc) expression, and into ten regions, named lobules, by morphological division along the anterior-to-posterior axis(*29*), here the expression frequency of cPcdhβ in Purkinje cells was examined in relation to Zebrin II expression and lobules. The expression frequency of β3-tdTomato in Purkinje cells was 2.4 ± 0.2% for the whole cerebellum. The β3-tdTomato reporter mice demonstrated stochastic tdTomato expression in both Zebrin II-positive and -negative Purkinje cells, and the frequency of tdTomato expression did not differ between Zebrin II-positive and -negative Purkinje cells, suggesting no detectable spatial bias in relation to Zebrin II expression (Figs. 2A-2B). In contrast, when comparing the expression frequency of β3-tdTomato between individual lobules, the mean values for individual lobules were similar, except for a minor decrease in lobule 7 and a minor increase in lobule 10 (Figs. 2C-2D). These results revealed that cPcdhβ provides single-cell identity with a weak spatial bias in relation to cerebellar lobules.

**Fig. 2.**
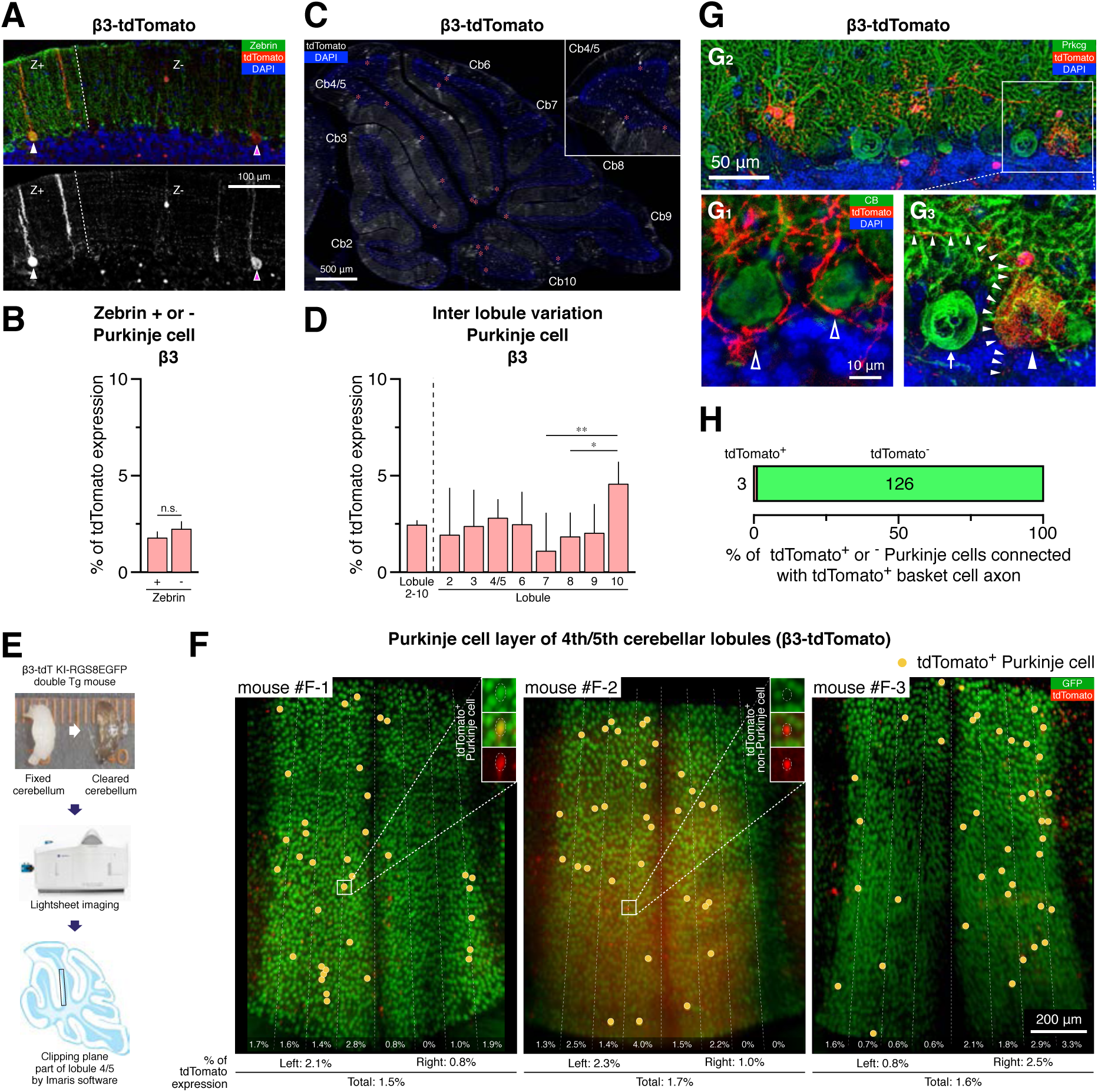
Spatial patterning of stochastic cPcdhβ3 expression in cerebellum. (A) Expression of cPcdhβ3 in both Zebrin II (+) and (-) Purkinje cells. (B) Quantification of the frequency of cPcdhβ3 (+) Purkinje cells in Zebrin II (+) and (-) Purkinje cells. There is no significant difference between the two groups. Data points, mean ± SD. n = more than 500 cells for each group. (C-D) Spatial biases of cPcdhβ3-positive frequencies in Purkinje cells among each cerebellar lobule. (E-F) Left-right asymmetry and inter-individual variation of cPcdhβ3 expression. (E) Schematic drawing of the method. (F) Expression patterns of tdTomato on Purkinje cells at the ventral part of the 4th/5th cerebellar lobule in three individuals (mouse #F-1, #F-2, and #F-3). Cell bodies of RGS8-GFP positive Purkinje cells are indicated by Green, tdTomato-positive cells are indicated by Red, and tdTomato- and RGS8-GFP double-positive Purkinje cells are marked by Yellow dots. Numbers at the lower part of the picture indicate the frequencies of tdTomato and RGS8-EGFP double-positive Purkinje cells among RGS8-positive Purkinje cells of each bin, left or right part, and entire photo, respectively. (G-H) Frequency of cPcdhβ3 expression is independent of connection with cPcdhβ3 (+) basket cells. (G) Representative image of tdTomato (-)(G1) and (+)(G2 and G3) Purkinje cells connected with tdTomato (+) basket cell axon. (H) Quantification of the frequency of tdTomato (-) and (+) Purkinje cells. There is no significant difference from the tdTomato (-) and (+) Purkinje cells in the whole cerebellum. Numbers indicate cells for each group.

Second, we investigated whether the spatial distribution of the cPcdhβ expression was identical between individual mice, the cPcdhβ expression at the same location in different mice were examined (Figs. 2E-2F). Cerebellar Purkinje cells can be reliably identified based on their monolayer location, size, and marker expression (in this study, we used Rgs8-EGFP transgene)(*30*). Purkinje cells on the ventral part of cerebellar lobules 4/5 showed a planar arrangement. This system allowed us to compare stochastic cPcdhβ expression between the same locations in different animals.

Three β3-tdTomato reporter mice from the same litter and of the same sex (all male) and postnatal age (day 9) were fixed, the cerebella cleared and imaged with a light-sheet microscope, and the vermis was 3D reconstructed. The ventral region of lobule 4/5 was then extracted from the reconstructed image, and the spatial distribution of tdTomato^+^ Purkinje cells was examined and compared between the three animals. The Rgs8-EGFP transgene expressed EGFP in most cerebellar Purkinje cells but not in the midline region of lobule 4/5(*30*, *31*). The distribution of tdTomato^+^ Purkinje cells, marked with yellow circles in Fig. 2F, showed inter-individual variation. Interestingly, the spatial distribution of cPcdhβ expression in Purkinje cells showed left-right asymmetry. Although the analysis is restricted to cerebellar Purkinje cells, these results provide strong evidence that the cPcdhβ expression follows a spatially-biased stochasticity.

### Connection is less correlated with cPcdhβ3 expression at basket cells to Purkinje cell synapse

A number of studies have shown that the homophilic interaction of cPcdh leads to repulsion between cell membranes with the same repertoire of cPcdh. However, the impact of cPcdh repertoire on synaptic connection between cells with the same cPcdh expression requires further investigation. To address this issue, we examined the correlation between cPcdhβ3 expression and synaptic connection in cerebellar basket cells that synapse with Purkinje cells. This neural circuit is advantageous because it allows easy visual identification of pre- and postsynaptic cells. This is because the basket cell axon extends in an anteroposterior axial direction and its branches surround the Purkinje cell body, resulting in synaptic connection.

Sagittal slices of the cerebellum from 21-day-old β3-tdTomato reporter mice, just after circuit maturation, were double immunostained for tdTomato and Purkinje cell markers (PKCγ or Calbindin)(Fig. 2G). We identified 55 tdTomato-positive basket cells and 129 Purkinje cells that were connected to them. The frequency of tdTomato-positive Purkinje cells was 2.3%, which is comparable to the frequency of tdTomato-positive Purkinje cells in the cerebellum (2.4 ± 0.2%, Fig. 2D). This suggests that cPcdhβ3 expression might have less influence on basket cell to Purkinje cell synaptic connection and that cPcdhβ3 expression follows a synaptic connection independent stochasticity. These results demonstrate that the cPcdh reporter mouse can improve our understanding of the relationship between synaptic connection and cPcdh expression.

### The frequency transition of cPcdhβ3 expression is different between cell types

We next examined the temporal aspects of stochastic cPcdhβ expression in the mouse brain. The qRT-PCR analysis for the expression of cPcdhα, -β, and -γ confirmed the previous results(*3*, *32*, *33*), and showed that all cPcdhα, -β, and -γ, including the modified cPcdhβ3, show comparable expression transitions (Fig. 3A). These results suggest that the cPcdh expression is dynamic in the brain, and the β3-tdTomato reporter mouse will be suitable for monitoring the dynamics of cPcdhβ3 expression at single-cell resolution.

**Fig. 3.**
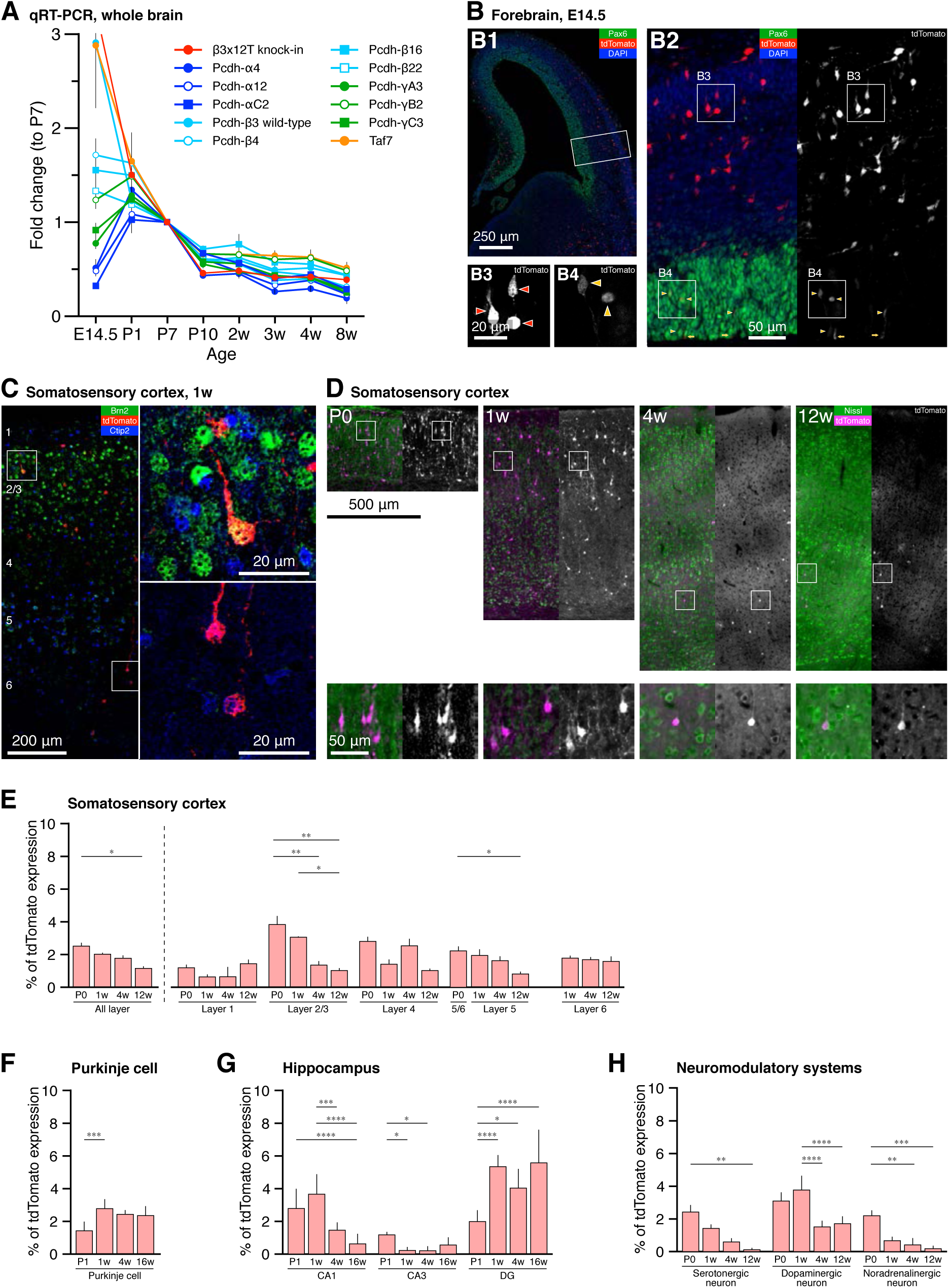
Stochastic cPcdhβ3 expression in developing brain. (A) The mRNA levels of cPcdhα, cPcdhβ, Pcdhγ genes and tdTomato were measured in the whole brain of β3-tdTomato mouse (heterozygous) by qRT-PCR (n = 3). The y-axis represents the fold change in mRNA levels relative to postnatal day 7 (P7). cPcdh expression decreased during maturation. (B)Expression of Pcdhβ3 in the embryonic forebrain. (B1) Lower magnification view of coronal sections of the embryonic forebrain from β3-tdTomato mice at E14.5. (B2) High-magnification image of the boxed area in (B1). (B3 and B4) High-magnification images of the boxed area in (B2). Both Pax6-positive cells (neural stem/progenitor cells) and Pax6-negative cells (post-mitotic neurons) stochastically expressed tdTomato. Yellow arrowheads: tdTomato and Pax6 double-positive cells. Red arrowheads: tdTomato-positive and Pax6-negative cells. (C)Coronal sections of the somatosensory cortex from β3-tdTomato mouse at 1-week-old. The boxed region is magnified at the right. (D)Expression of cPcdhβ3 in the developing somatosensory cortex at postnatal days 0, 1-week-old, 4-week-old, and 12-week-old mice. The boxed region is magnified at the bottom of the figure. (E-H) Quantification of cPcdhβ3 expression frequency during brain maturation at postnatal day 0 (P0), 1 week (1w), 4 weeks (4w), and 12 or 16 weeks (12w or 16w). Data points are presented as the mean ± SE. n = 2-3. (E) Somatosensory cortex (left) and laminar distribution patterns (right), (F) Purkinje cells, (G) hippocampus, and (H) neuromodulatory neurons.

Thus, we examined cPcdhβ3 expression dynamics in the cerebral cortex, where the birthdate, birthplace, migration, and final destination of most neurons and glia are well documented(*34*). Histological analysis of the β3-tdTomato embryonic telencephalon revealed that a subset of neural stem/progenitor cells express cPcdhβ3 and its expression level was weaker than that in post-mitotic neurons (Fig. 3B), consistent with the previous finding(*35*). Stochastic cPcdhβ3 expression in neural stem/progenitor cells begins as early as E12.5 (data not shown). The spatial distribution of cPcdhβ3^+^ cells along the medial-to-lateral axis in the postnatal cerebral cortex does not seem to have a columnar organization, rather a salt-and-pepper organization along the medial-to-lateral axis (Fig. 3C-3D).

The frequency of cPcdhβ3^+^ cells showed dependency on the cortical layers: the frequency in layer 2/3 was the highest at postnatal day 0, whereas that in layer 4 was the highest at 1 week (Fig. 4D and 4E). Interestingly, the frequency of cPcdhβ3^+^ cells in a given layer differed between developmental timings: layer 4 showed the highest frequency at 4 weeks of age, while the other layers had the highest frequency at postnatal day 0. Since neuronal cell bodies almost settle in the postnatal cortex, these observations suggest that the cPcdhβ3 expression changes over time.

**Fig. 4.**
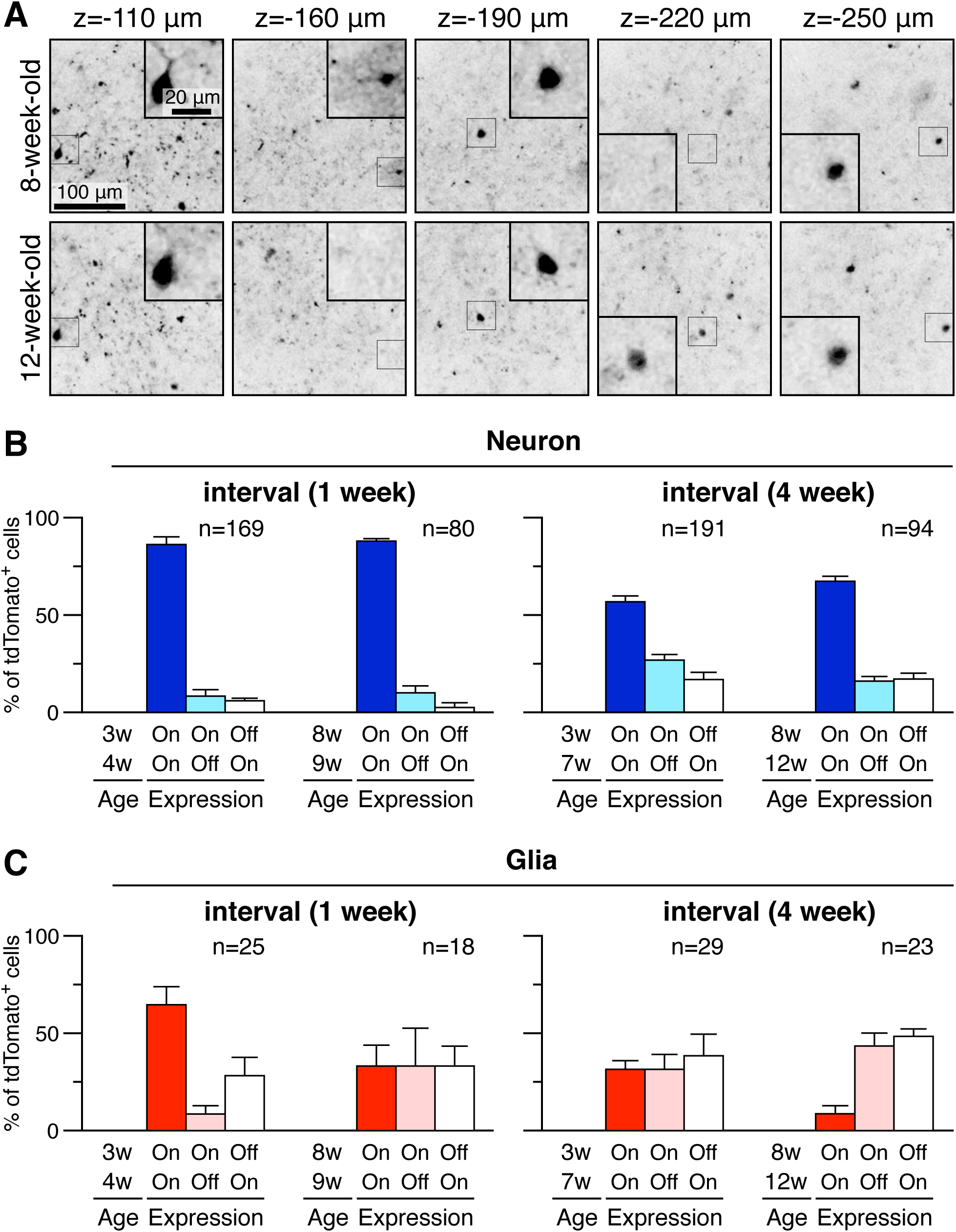
Stochastic and dynamic cPcdhβ3 expression in living brain of adult mouse. Expression of cPcdhβ3 was assessed using in vivo two-photon microscopy in the somatosensory cortex of β3-tdTomato reporter mice at the juvenile and young adult stages, and Pcdhβ3 expression in the same regions was determined at 1 week and 4 weeks later. (A) Representative two-photon laser microscopy images of the living somatosensory cortex from β3-tdTomato reporter mouse at 8 weeks (upper panels) and 12 weeks (lower panels). Black: β3-tdTomato^+^ cells. The inset panels show higher-magnification images of the boxed areas. The depth from the surface of the brain is indicated in the upper part of the pictures. Examples of a neuron whose β3-tdTomato expression is unchanged between the two time points are shown at depths of z=-110, -190, and -250 µm. An example of a glial cell switching β3-tdTomato expression from ON to OFF is shown at a depth of z=-160 µm. An example of a neuron switching β3-tdTomato expression from OFF to ON is shown at a depth of z=-220 µm. Scale bars: 100 and 20 µm. (B) Quantification of the number of neurons with unchanged expression at both ages (blue), switched expression from ON to OFF (aqua), and from OFF to ON (white). (C) Quantification of the number of glial cells with unchanged expression at both ages (red), expression switched from ON to OFF (pink), and from OFF to ON (white). Data obtained at 1-week intervals are shown in the left panel, and at 4-week intervals are shown in the right panel, respectively. Numbers represent the number of cells analyzed.

The changes in the frequency of cPcdhβ3^+^ cells during postnatal development were also consistent with other neuron types, including cerebellar Purkinje cells, hippocampal neurons in the CA1, CA3, and DG layers, and monoaminergic neurons (dopaminergic neurons in the VTA, serotonergic neurons in the dorsal raphe nucleus, and noradrenergic neurons in the LC) (Fig. 3F-3H and Fig. S5). Taken together, cPcdhβ expression is dynamically regulated throughout development, and its transition differs between cell types.

### Dynamics of the cPcdhβ3 expression in the living mouse brain

The previous section suggested that the cPcdhβ3 expression is developmentally dynamic in the brain. This finding raises the question of whether cPcdhβ expression at the single-cell level showed only decreased/turned-off or whether it is also increased/turned-on. We observed the cerebral cortex of a live β3-tdTomato mouse using 2-photon microscopy to detect the cPcdhβ3-expressing cells. The photographs in Fig. 4A are representative images of stochastic cPcdhβ3 expression at the same site in the somatosensory cortex of a living mouse at depths of ∼110 μm to ∼250 μm at 8-week-old (upper panels) and 12-week-old (lower panels). We identified identical cells based on their relative location to blood vessels and their dendritic morphology. The leftmost, middle, and rightmost neurons express cPcdhβ3 at 8- and 12-week-old. In contrast, one cPcdhβ3-expressing cell was observed in the enclosed area at ∼160 μm depth at 8 weeks, but no β3-expressing cells were observed at the same position at 12 weeks (left-second). Interestingly, while no β3-positive cells were observed in the enclosed area at a depth of ∼220 μm depth at 8 weeks, one cell expressed tdTomato in the same area at 12 weeks. Quantification of the images showed that approximately 90% of the neurons expressed tdTomato after a 1-week interval, while approximately 5% of neurons turned-off the tdTomato expression, while another 5 % of neurons turned-on the tdTomato expression (Fig. 4B). These rates were comparable between the 3–4-week-old and 8–9-week-old mice. Surprisingly, when compared with 1-week interval, there was an increase in the number of cells with altered expression at 4-week intervals. Fig. S6 shows that changes in cPcdhβ3 expression also occur in inhibitory neurons and that cPcdhβ3 expression level is altered in a living cell rather than alteration by cell death or cell migration. These results indicate that cPcdhβ3 expressions in neurons changed over time, and the changing rate was approximately 10% of neurons per week.

Next, we examined temporal changes in the expression of cPcdhβ3 in glial cells (Fig. 4C). The expression of cPcdhβ3 in glial cells changed over time faster than in neurons, ∼30% per week. These results reveal that cPcdhβ3 expression is dynamic at the single-cell level in the postnatal cerebral cortex and that its expression is not only turned-off but also turned-on, even at the adult stage.

### Modification of the endogenous cPcdhβ19 and visualization of monoallelic Pcdhβ19 expression *in vivo*

To further extend the cPcdhβ reporter mouse regarding color and isoform, we generated two additional lines of knock-in mice. One is knocked in an EGFP cDNA into the endogenous cPcdhβ19 locus, and the other is knocked in a multi-tandem tdTomato into the endogenous cPcdhβ19 locus (Fig. 5A, Fig. S7). We selected the cPcdhβ19 locus because it has been demonstrated that cPcdhβ3 and cPcdhβ19 genes are regulated by distinct mechanisms(*36*). Both the heterozygous and homozygous knock-in mice were fertile and did not exhibit any morphological or developmental defects.

**Fig. 5.**
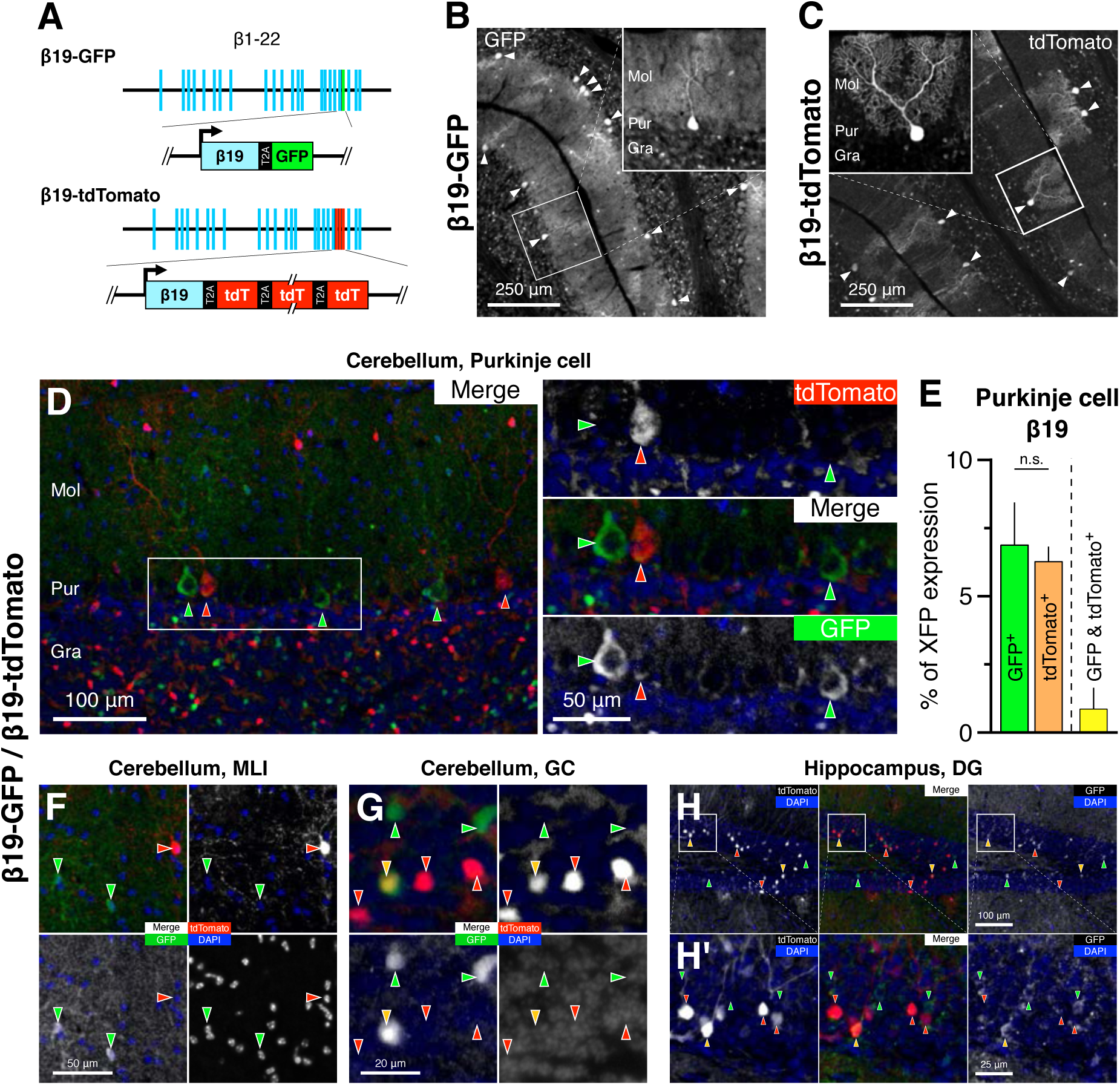
Generation of β19-tdTomato reporter mouse and visualization of monoallelic cPcdhβ19 expression in vivo. (A) Genomic structure of β19-GFP and β19-tdTomato reporter alleles, which includes single GFP or multi-tandem tdTomato in the endogenous cPcdhβ19 locus, respectively. (B) GFP expression from the Pcdhβ19-GFP reporter allele (4-week-old) in the cerebellum sagittal section. (C) tdTomato expression from the Pcdhβ19-tdTomato reporter allele (4-week-old) in the cerebellum sagittal section. (D-H) Visualization of monoallelic cPcdhβ19 expression in vivo. Sections of double Tg mouse harboring β19-GFP and β19-tdTomato reporter alleles shows monoallelic cPcdhβ19 expression in cerebellar Purkinje cells (D), molecular layer interneuron (F), granule cell (G), and granule cells in hippocampal dentate gyrus (H). Quantification of monoallelic or biallelic cPcdhβ19 expression in cerebellar Purkinje cells (E). n=3.

We examined the β19-GFP and the β19-tdTomato reporter mice using histological analysis to confirm that the reporter genes recapitulated the endogenous expression of cPcdhβ19. The β19-GFP and β19-tdTomato mice exhibited stochastic GFP and tdTomato expression in the cerebellum, respectively (Fig. 5B and 5C), which closely resembled the native cPcdhβ19 expression (Fig. S1F). The frequency of GFP^+^ or tdTomato^+^ Purkinje cells is comparable to our previous data obtained by single-cell RT-PCR: 7.5 ± 0.3% for β19-tdTomato, 6.9 ± 1.5% for β19-GFP, and 10.7% for endogenous cPcdhβ19 (Fig. 5E) (*15*). It is crucial to note that the frequencies of XFP^+^ cells observed in the β19-GFP or tdTomato reporter mice were significantly higher than those observed in the β3-tdTomato mice. This recapitulates the endogenous expression of cPcdhβ3 and cPcdhβ19(*15*), validating the β19-GFP and the β19-tdTomato reporter mice.

While immunostaining using anti-GFP antibody is necessary for reliable GFP detection in β19-GFP mouse brain, the tdTomato fluorescence can be detected without immunofluorescence enhancement in β19-tdTomato mouse brain. The β19-tdTomato mouse is mainly used for the expression analysis of cPcdhβ19 *in vivo*.

In the β19-tdTomato reporter mice, tdTomato is expressed in a subset of neurons throughout nervous systems, including the retina, olfactory bulb, cerebral cortex, PV^+^ neurons in the cerebral cortex, caudate putamen, calbindin^+^ neurons in the inferior olivary nucleus, serotonergic neurons in the dorsal raphe, dopaminergic neurons in the ventral tegmental area, noradrenergic neurons in locus coeruleus, granule cell in the cerebellum, astrocytes in the cerebral cortex, and Bergmann glia in the cerebellum, suggesting that the Pcdhβ19 expression is stochastic in these cell types (Fig. S8). In contrast, the photographs showed no tdTomato expression in microglia, blood vessels, testis, nor kidney suggesting that Pcdhβ19 is not expressed in these cells.

We next examined the spatial distribution of Pcdhβ19 expression in cerebellar Purkinje cells. We found that the frequency of tdTomato expression did not differ between Zebrin II-positive and -negative Purkinje cells in β19-tdTomato reporter mice, suggesting no detectable bias in relation to Zebrin expression (Fig. S9A-B). In contrast, when comparing the expression frequency of β19-tdTomato between individual lobules, a minor decrease in lobule 7 was observed. These results showed similar trends with the β3-tdTomato reporter mice, thus suggest that Pcdhβ provides single-cell identity with a weak spatial bias in cerebellar lobules.

We examined Pcdhβ19 expression in the developing cerebral cortex. First, we observed the cerebral cortex of embryonic β19-tdTomato mice. As in the case of b3-tdTomato, expression was low in neural stem/progenitor cells, while expression was high in postmitotic neurons. The layer-by-layer expression frequency varied according to developmental timing from P0 to 4-week-old. Second, to examine whether the left-right asymmetric distribution of the Pcdhβ19 expression can be observed as similar to Pcdhβ3, we examined the spatial distribution of tdTomato-positive cells in the cerebral cortex using β19-tdTomato reporter mouse. The distribution of tdTomato-positive cells beneath the midline demonstrated a left-right asymmetry (Fig. S9E-F). Third, to analyze the relationship between Pcdhβ expression and cell lineage, we crossed β19-tdTomato mice with TFC.09 × loxP Ai32 reporter mice and histologically analyzed tdTomato and YFP expression in the cerebral cortex (Fig. S10). The results revealed that only a subset of YFP-positive neurons and astrocytes expressed tdTomato, indicating that Pcdhβ expression in a given cell lineage is not homogeneous, at least in cortical excitatory neurons and glia. These results demonstrate that the β19-tdTomato reporter mouse are valuable for investigating the spatiotemporal distributions of cPcdhβ19 expression *in vivo*.

Finally, we examined monoallelic cPcdhβ expression by combining the β19-GFP and the β19-tdTomato reporter mouse lines (Fig. 5D-H). Although previous studies have revealed that monoallelic cPcdh expression diversifies the cPcdh single-cell identity, only Purkinje cells have been demonstrated to show monoallelic cPcdh expression. Cerebellar sections from double transgenic mice harboring β19-tdTomato and β19-GFP alleles showed distinct labeling between tdTomato^+^ and GFP^+^ Purkinje cells, providing direct evidence of the monoallelic expression of cPcdhβ19 from these knock-in alleles (Fig. 5D and 5E). The cerebellar and hippocampal sections from double transgenic mice harboring β19-tdTomato and β19-GFP alleles showed distinct labeling between tdTomato+ and GFP+ in the molecular layer interneurons and granule cells in the cerebellum and hippocampal dentate gyrus, providing direct evidence of monoallelic expression of cPcdhβ19 in these neurons, expanding the cell types showing monoallelic cPcdh expression.

### Brain dysfunction leads to dysregulation of stochastic cPcdhβ expression in cell-type-dependent manner

Dysregulation of the cPcdh genes is associated with brain dysfunction(*5*, *9*, *21*, *37–41*). However, the spatiotemporal distribution and cell-type specificity of the dysregulation in the cPcdh cluster remain unknown. In this study, we examined the expression of cPcdhβ3 and cPcdhβ19 in intellectual disability model mice and β3 expression in the aging brain.

At first, we examined the expression of cPcdhβ3 and cPcdhβ19 in the brains of the Ehmt1 heterozygous deficient Kleefstra syndrome (KS) mouse model, which shows intellectual disabilities and autistic behavior similar to that in humans(*42*). Recently, Iacono et al. revealed that the expression levels of cPcdh genes are downregulated in the KS mouse hippocampus(*38*). However, the spatiotemporal distribution and cell-type specificity of cPcdh dysregulation in KS mouse models remain unclear. Therefore, we examined the expression of cPcdhβ3 and cPcdhβ19 in the hippocampus and cerebellum of KS model mice using the β3- and β19-tdTomato reporter mice (Fig. 6A and S6). The frequency of β3- and β19-expressing cells was different between wild-type and KS model mice, and interestingly, it showed cell-type dependency. In contrast, it was not altered in the cerebellar Purkinje cells. Interestingly, the tdTomato fluorescence at a single cell was comparable between the wild-type and KS model mice, suggesting a comparable expression level in a single cell. These results revealed that the downregulation of cPcdh expression in KS model mice is the sum of cell-type-dependent alteration of expression frequency and maintenance of expression level at a single-cell level.

**Fig. 6.**
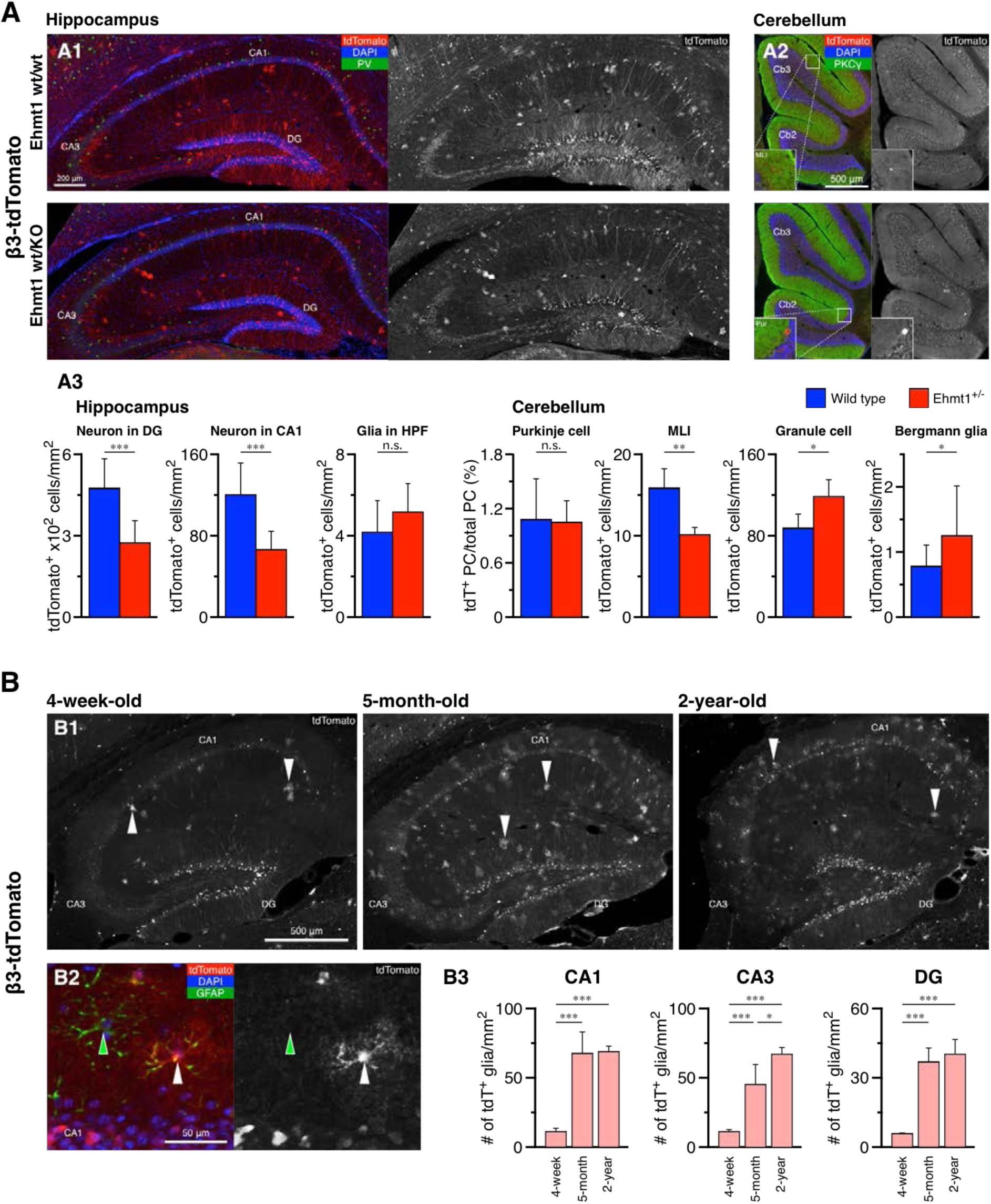
Stochastic cPcdhβ3 expression under pathological and aged condition. (A) Decreased expression of cPcdhβ3 in intellectual disability model mouse. Immunohistochemical staining of the hippocampus (A1) and cerebellum (A2) in control and Ehmt1^+/-^ mouse at 4 weeks. (A3) Quantification of tdTomato-positive cells in control (blue) and Ehmt1^+/-^ mouse (red). n = 3. (B) Increasing cPcdhβ3 positive glia cells in aging mouse brain. (B1) Representative images of β3-tdTomato mouse hippocampus at 4 weeks (left), 5 months (middle), and 2 years (right). (B2) Stochastic tdTomato expression in astrocyte of the hippocampus (5-month-old). (B3) Quantification of tdTomato^+^ glia cells in the hippocampus. n = 3.

Next, we examined the cPcdhβ3 expressions in the aging brain using β3-tdTomato reporter mice. Our results clearly show that the number of tdTomato-positive neurons was mildly reduced in the aged brain, whereas a significant increase in tdTomato-positive glial cell density was observed in the hippocampus (Fig. 6B). These findings demonstrate that the dysregulation of cPcdh genes upon aging is cell-type-dependent. These results clearly demonstrate that alterations in cPcdh expression in brain dysfunction are cell-type-dependent. This highlights the crucial importance of single-cell-resolution analysis for fully understanding the impact of brain dysfunction on cPcdh cellular barcode.

## Discussion

The difficulty in examining spatiotemporal patterning of stochastic cPcdh expression has hindered our understanding of the physiological roles mediated by cPcdh cellular barcode. Furthermore, cPcdh dysregulation has been reported in several pathological conditions, but examination at single-cell resolution is yet to be conducted. This is due to a lack of reliable method for detecting each single cPcdh isoform. We have overcome this by developing several lines of XFP (tdTomato or GFP) knock-in mice. This has enabled us to assess the expression of cPcdhβ3 or cPcdhβ19 in fixed and living brains from embryonic to aged stage. Our cPcdhβ reporter mice are therefore the ideal model for studying spatiotemporal patterning, physiological roles and pathological contributions of cPcdh cellular barcode *in vivo*.

### Advantage of the cPcdhβ reporter mouse lines

This study employed a genetic approach to map the spatiotemporal patterning of cells expressing a single cPcdhβ gene in the mouse brain. Our reporter mice are significantly more specific than previously reported methods, including immunofluorescence, mRNA *in situ* hybridization, or image-based *in situ* transcriptomics methods (STARmap, MERFISH, ISS, and BOLORAMIS), in detecting cells expressing a single cPcdhβ gene(*43–46*). Our reporter mice require less laborious processing and can be used to identify cell types in intact tissue and living brains. Another advantage is that our mice can report translation-level expression with XFP.

### Cell-type and spatial dependent regulation of stochastic cPcdhβ expression

This study revealed that the frequency of cPcdhβ expression is influenced by the cell type and developmental stage. These results suggest that the population-level cPcdh expression is regulated by a deterministic manner. In contrast, expression at the single-cell level is stochastically regulated. Thus, the cPcdh cellular barcode was constructed using deterministic and stochastic regulatory modes. The frequency of *Drosophila* Dscam1 expression, which is responsible for the generation of cellular barcode in the invertebrate nervous system, also differs between cell types, indicating the presence of a deterministic regulatory mechanism(*47*). Further, alternative splicing, not promoter choice, is known to regulate Dscam1 expression(*48*). Thus, the collaboration of dual gene regulatory modes, including deterministic and stochastic regulation, may have evolved as a common strategy, albeit via different molecular mechanisms by which neurons acquire single-cell identities.

Analysis of cPcdhβ3 and cPcdhβ19 expression in cerebellar Purkinje cells revealed a spatial bias in their expression frequency along the rostral-to-caudal axis (Figs. 2C and 2D). This suggests that promoter choice in the cPcdhβ cluster is related to spatial information in the brain, at least in the cerebellar Purkinje cells. It has been reported that cPcdh expression is regulated by numerous transcription factors and DNA-binding proteins, including CTCF, NRSF/REST, Sox4, Egr-1, Nipbl, Mecp2, Dnmt3b, Ehmt1, Wiz, Suv39h2, Smchd1, Mpp8, Morc2, HUSH complex, Setdb1, and WAPL (*9*, *41*, *49–54*). Additionally, cPcdh promoter CpG methylation is inversely correlated with cPcdh expression(*12*, *27*, *55*). Although the critical determinant of cPcdhβ expression in Purkinje cells remains unknown, some of these factors may be distributed in a spatially biased manner.

We noted that the dysregulation of cPcdhβ expression by intellectual disability or aging differs between neurons and glia (Fig. 6 and s6), suggesting that dysregulation of cPcdh expression is cell-type-dependent. cPcdh has previously been shown to affect dendrite or axon self-avoidance, dendrite arborization, spatial arrangement of the cell body, synaptogenesis, and astrocyte-neuron contact(*4*, *5*, *19*, *56*). Alterations in histone modification or DNA methylation in the cPcdh locus have been demonstrated in intellectual disability and aging(*37*, *57–60*). These observations indicate that intellectual disability or aging influence these characteristics with cell-type-dependency. Future studies using human patient samples and other disease models are needed to test the potential pathogenic mechanisms inferred in this study.

We also examined the cPcdhβ expression in non-neural cells (Fig. S4O and S4P). Kidney section from β3- and β19-tdTomato reporter mice showed no tdTomato expression. Surprisingly, the testis sections from β19-tdTomato reporter mice showed no expression of tdTomato, whereas the sections from β3-tdTomato reporter mice showed constitutive expression of tdTomato in differentiated sperm(*28*), clearly demonstrating the gene regulatory difference between β3 and β19 in the testis. Furthermore, the differentiated sperm in the testis is interesting because it expresses β3-tdTomato constitutively, not stochastically, indicating stochastic cPcdhβ3 expression requires specific gene regulatory mechanisms that are absent in the testis.

### Dynamics of cPcdhβ expression *in vivo*

This study revealed the cPcdhβ expression dynamics *in vivo*. The changes in the frequency of cPcdhβ3^+^ cells during postnatal development were consistent with several neuron types, including cerebellar Purkinje cells, hippocampal neurons in the CA1, CA3, and DG layers, and monoaminergic neurons (dopaminergic neurons in the VTA, serotonergic neurons in the dorsal raphe nucleus, and noradrenergic neurons in the LC) (Fig. 3). Although previous reports have shown very weak expression in adult serotonergic neurons(*20*), expression of cPcdhβ3 and cPcdhβ19 was detected at P0 (Fig. 3 and data not shown). These results suggest that stochastic cPcdhβ expression is dynamically regulated throughout development, and its transition differs between cell types.

Live imaging of β3-tdTomato reporter mice showed that cPcdhβ expression changed over time in a single cell in the cerebral cortex. These changes include not only disappearances but also new appearances. These results suggest that cPcdh cellular barcode can change dynamically in the brain of living mice. It should be noted that the timing of expression changes differs among neurons located close to each other, suggesting that expression changes are not constrained to a particular brain region. Although its physiological significance requires further examinations, the dynamic cPcdhβ expression could minimize the risk of neighboring neurons/glia sharing the same cPcdh isoforms over an extended period.

The underlying molecular mechanisms of this dynamic cPcdhβ expression is interesting. Although the half-life of cPcdhβ mRNA is unknown, mRNA generally has a mean half-life of ∼7 hrs(*61*). Therefore, changes in cPcdhβ expression at the several-week time scale are likely due to contributions other than mRNA degradation and possibly changes in cPcdhβ promoter choice. cPcdhβ promoter choice has been suggested to be influenced by DNA methylation, histone modifications, chromatin structure, and transcription factors(*5*, *9*, *21*, *38*, *50*, *62*). However, the appearance of cPcdhβ3-expressing cells makes gene silencing mediated by DNA methylation and/or constitutive heterochromatin unlikely.

In addition to cPcdh, at least three systems utilize a stochastic mechanism to diversify cell surface molecule expression in the animal kingdom, viz., the mammalian immunoglobulin (IgG), mammalian olfactory receptors (OR), and *Drosophila* Dscam1(*2*). Among these, the expression of IgG and OR are stable once the expression is established; however, the expression of Dscam1 are dynamic(*47*). Dscam1 also provides a single-cell identity in the *Drosophila* nervous system. Thus, it suggests that the expression dynamics of genes that provide a single-cell identity in nervous system may play special yet unknown physiological roles in the central nervous system.

## Conclusion

Our findings consistently support the utility of the new cPcdhβ reporter mouse lines for histological and functional studies. These models are of particular interest for studies of memory, psychiatric disorder andneuronal survival, since cPcdhβs play a role in these situations(*38*, *63*, *64*). cPcdhβs is evolutionarily conserved, including humans(*7*), suggesting a translational utility of rodent studies. The cPcdhβ reporter mouse lines offer a tool to examine the functional role of cPcdh cellular barcode *in vivo*.

## Acknowledgments

We thank Drs. A. Oue, N. Sugo, H. Kobayashi, E. Tarusawa, and members of the Yanagawa and Yagi laboratories for their helpful discussions. We acknowledge Mss. R. Natsume, T. Kakinuma, S. Sato, W. Mizutani, Y. Morimoto, and M. Higuchi for providing technical support. We thank the members of the Bioresource Center, Gunma University, and the Animal Facility, Osaka University Frontier Biosciences, for maintaining the animals used in this study. We greatly appreciate Dr. Neal Copeland for providing the BAC recombinant reagents, Dr. Roger Tsien for providing the tdTomato cDNA, Dr. Callos Lois for providing the TFC.09 mouse, and The GENSAT project for providing the Rgs8-EGFP mice.

## Funding

JSPS KAKENHI [Grant-in-Aid for Scientific Research (C)] 24500375, 15K06696, 22K06457 (RK)

JSPS KAKENHI [Fund for the Promotion of Joint International Research (Fostering Joint International Research)], Grant number: 15KK0331 (RK)

JSPS KAKENHI [Grant-in-Aid for Scientific Research on Innovative Areas “Integrative research toward elucidation of generative brain systems for individuality”] 17H05937, 17H05967, 19H04895, 19H04922 (RK, YIU)

JSPS KAKENHI [Grant-in-Aid for Scientific Research on Transformative Research Areas (A) Adaptive Circuit Census] 22H05498 (TY)

JSPS KAKENHI JP 16H06276 (AdAMS)

JSPS KAKENHI JP16H06280 (AbiS)

Planned Collaborative Project, and the Cooperative Study Program of the National Institute for Physiological Sciences (RK, TY)

## Author contributions

Conceptualization: RK, TY

Methodology: RK, MA, YIU, YM, TI, MW, KS, YY

Investigation: RK, YT, SH, AM

Visualization: RK, YT

Funding acquisition: RK, YIU, TY

Project administration: RK, TY

Supervision: TY

Writing – original draft: RK, TY

Writing – review & editing: RK, YT, MA, YIU, SH, TI, MW, KS, YY, TY

