## Supplemental file for "Visualization of stochastic expression of clustered protocadherin β isoforms *in vivo*"

**This PDF file includes:**

Materials and Methods

Figs. S1 to S11

### Materials and Methods

#### Mice

C57BL/6J (B6) and ICR mice were purchased from Charles River (Japan). The β3-tdTomato, β19-GFP mice, and β19-tdTomato mice were generated and maintained on a B6 background (as shown below). The Pcdhab$^{del/del}$ mouse (Hasegawa et al., 2016) was maintained on a B6 background. The Rgs8-EGFP mouse (STOCK Tg(Rgs8-EGFP)CB132Gsat/Mmucd) was a generous gift from the GENSAT project and maintained on a B6 background (Kaneko et al., 2018). The TFC.09 mouse was a generous gift from Dr. Carlos Lois and was maintained on a B6 background.

Mice were group-housed (a maximum of five mice per cage) and maintained on a 12 hr light–dark cycle (8 am to 8 pm, light), with access to food and water ad libitum. Male and female mice were used for the morphological experiments. Embryonic stages were calculated by defining noon of the day of vaginal plug observation as embryonic day 0.5 (E0.5) and the day of birth as postnatal day 0 (P0). All experiments involving animals were approved by our institute's Animal Care and Use Committee and conformed to Japanese guidelines.

#### Generation of mutant mice

All targeting vectors were constructed using standard molecular cloning approaches.

To generate cPcdhβ3-tdTomato knock-in mouse (β3-tdTomato reporter mouse), we constructed a targeting vector harboring the 12-tandemly arrayed tdTomato flanked with T2A self-cleaving peptide downstream of the C-terminally Myc tagged cPcdhβ3 gene (without a stop sequence) and an FRT-flanked neomycin resistance cassette (neo-cassette) from neo plasmid. The targeting vector was introduced into C57BL/6N ES RENKA cells by electroporation as described previously(*65*). After isolating homologous recombinants in ES cells, the clones were introduced into 8-cell embryos of ICR mice. To delete the neo-cassette, chimeric mice were mated with CAG-FLPe mice (C57BL/6-Tg(CAG-flpe)36 (B6-CAG-flpe), a generous gift from Itohara S, RIKEN BRC stock no. RBRC01834) that ubiquitously expresses FLPe recombinase. For experiments using heterozygous β3-tdTomato reporter mouse, we used mice generated by crossing homozygous β3-tdTomato reporter males with C57BL/6 females.

To generate cPcdhβ19-GFP knock-in mice (β19-GFP reporter mice), we constructed a targeting vector harboring EGFP flanked by the T2A self-cleaving peptide downstream of the C-terminally PA tagged cPcdhβ19 gene (without a stop sequence). The targeting vector was introduced into one-cell-stage zygotes, which were obtained by mating B6C3F1 males with superovulated females by microinjection, as described previously(*66*). Correctly targeted pups were identified using genomic PCR. For experiments using heterozygous β19-GFP reporter mice, we used mice generated by crossing homozygous β19-GFP reporter males with C57BL/6 females.

To generate cPcdhβ19-tdTomato knock-in mice (β19-tdTomato reporter mouse), we constructed a targeting vector harboring the 12-tandemly arrayed tdTomato flanked by T2A self-cleaving peptide downstream of the C-terminally PA tagged cPcdhβ19 gene (without stop sequence). The targeting vector was introduced into one-cell-stage zygotes, which were obtained by mating B6C3F1 males with superovulated females by microinjection, as described previously(*66*). Correctly targeted pups were identified using genomic PCR. For experiments using heterozygous β19-tdTomato reporter mice, we used mice generated by crossing homozygous β19-tdTomato reporter males with C57BL/6 females. For experiments using double transgenic mice harboring β19-GFP reporter and β19-tdTomato reporter alleles, we used mice generated by crossing homozygous β19-GFP reporter males with homozygous β19-tdTomato reporter females.

Histology and antibodies

Immunofluorescence staining was performed as described previously(*67*). Briefly, mice were anesthetized and perfused transcardially with PBS and 4% paraformaldehyde in PBS under deep anesthesia with ketamine/xylazine. Tissues (brains, eyes, kidneys, and testis) were removed, post-fixed (6 hr), and cryoprotected (25% sucrose for 3 d). Slices were prepared at 20-μm thickness using a cryostat (Leica).

For immunostaining, sections were blocked in 0.2% Triton X-100, 5% goat serum, and 2% bovine serum albumin in PBS and then incubated with diluted primary antibodies for 24-48 h at 4°C. After extensive washing in PBS, sections were incubated with Alexa dye–conjugated secondary antibodies for 1 hours before being washed again and mounted with CC/Mount mounting medium (DBS Diagnostic BioSystems). For staining with mouse monoclonal antibodies, the sections were blocked with monovalent Fab fragments (Jackson ImmunoResearch) to reduce background signals in the mouse tissues. To improve the visualization of brain regions, neurons were sometimes labeled with Neurotrace dyes (Invitrogen) or sections were counterstained with DAPI (Sigma-Aldrich). Sections were stored at -20°C until they were imaged.

The primary antibodies used were as follows: Aldolase C (guinea pig, Frontier Institute, 1/200), Calbindin (mouse, Swant, 1/1000), DCX (mouse, Santa Cruz, 1/50), GABA (rabbit, Sigma, 1/500), GFP (rabbit, Frontier Institute, 1/500), GST-π (rabbit, MBL, 1/400), Iba1 (rabbit, Fujifilm WAKO, 1/500), Ki67 (rabbit, Leica, 1/100), Parvalbumin (mouse, Swant, 1/1000), Pax2 (rabbit, Invitrogen, 1/100), Pax6 (rabbit, Millipore, 1/500), S100b (rabbit, Frontier Institute, 1/200), somatostatin (rat, Merck, 1/100), tdTomato (guinea pig, AB_2631185, 1/200), tdTomato (rabbit, AB_2571847, 1/200), Tbr2 (rabbit, ab23345, 1/500), TH (chicken, ab76442, 1/500), Tph2 (rabbit, Novus, 1/500), VAChT (rabbit, Frontier Institute, 1/200).

*In situ* hybridization (ISH) was performed essentially as described previously on 10-μm-thick frozen sagittal sections prepared from wild-type mice (4-week-old) using digoxigenin (DIG)-labeled cRNA probes. The probes for the cPcdh genes were the same as in our previous studies(*67*).

Images were acquired using a BZ-9000, BZ-X710, or BZ-X810 fluorescence microscope (Keyence), or Stellaris 5, or Stellaris 8 confocal microscope (Leica). Images were post processed with Fiji or Pixelmator (Pixelmator Team). The number of cells was counted using the Fiji.

Brain dissociation, cell sorting using a FACS

The brain dissociation protocol was performed with modification as previously described (Kaneko R et al, 2014). β3-tdTomato knock-in mouse homozygous female was crossed with Pcdhab$^{del/del}$ male, and embryonic mouse brains were harvested at E16.5 and placed in chilled PBS. The cortexes were micro-dissected in chilled PBS. The recovered cortexes were minced into smaller blocks and immediately transferred into pre-warmed dissociation medium (10 U papain [Worthington, Ohio, United States], and 2 mg DNase I [Sigma-Aldrich, Missouri, United States] in PBS. After a 30-min incubation at 37°C, tissues were gently dissociated using a fire-polished glass pipette (approximately 40 μm of internal diameter). Brain suspension was resuspended with PBS, followed by filtration using a cell strainer (pore size 40 μm). The filtrate was subjected to centrifugation, and the collected pellet was resuspended in 2% FBS in PBS. All procedures were conducted on ice except for the digestion process. The tdTomato-positive and -negative cells were sorted using a FACS (SONY, SH800).

DNA methylation analysis

The DNA methylation analysis protocol was performed with modification as previously described (Kaneko R et al, 2014). The genomic DNA was prepared with the EpiTect Plus LyseAll Kit (Qiagen) from FACS-sorted tdTomato-positive and -negative cells according to the supplier's recommendations. Bisulfite conversion was performed with the EpiTect Plus DNA Bisulfite Kit (Qiagen). PCR and DNA sequencing were carried out the same as previously described (Kaneko R et al, 2014).

Brain clearing and thick specimen imaging

For imaging of the cleared cerebellum, we cleared the cerebellum with SeeDB2 reagent and imaged it using light-sheet microscopy. Three male β3-tdTomato reporter and Rgs8-EGFP double-Tg mice from the same litter on postnatal day 9 were used. The PFA-fixed cerebellum was rinsed with PBS and cleared with SeeDB2G[74,75]. The cleared cerebellum was imaged using a light-sheet microscope (Zeiss Lightsheet Z.1, 25x objective). The images were 3D reconstructed using Imaris software, and the Purkinje cell layer of the ventral lobule 4/5 was extracted.

To image the cleared cerebral cortex, we cleared a thick section of the cerebral cortex with SeeDB2 reagent and imaged it using confocal microscopy. A β19-tdTomato reporter mouse was PFA-fixed on postnatal day 9. The cerebral cortex was then sliced using a cryostat (Leica C3050) at 0.3 mm thickness, rinsed in PBS, embedded in 10% fish gelatin, stained with DAPI then cleared with SeeDB2G. The cleaned samples were mounted on glass slides with a 0.3 mm thick plastic spacer. Samples were imaged using an Andor Dragonfly spinning disk confocal microscope (Oxford Instruments, UK) equipped with a 20x objective (Nikon). Individual image tiles were stitched into composite images using a Fusion Stitcher (Andor).

In vivo 2-photon imaging

In vivo 2-photon images were obtained using a thin-skull imaging window following a previously reported method(*67*, *70*) with minor modifications. Briefly, animals were anesthetized using a mixture of ketamine (100 mg/kg) and xylazine (10 mg/kg) at 1% ml/g body weight. The skull of each mouse was carefully thinned using a high-speed drill (Narishige) and a microsurgical blade (Surgistar no. 6400) after mice were anesthetized with an injection of ketamine/xylazine (22.5 mg/ml of ketamine and 1 mg/ml of xylazine in 0.9% NaCl i.p.; 5 ml/kg during the surgeries). Images were obtained using a 2-photon laser microscope customized for in vivo imaging (FVMPE-RS, Olympus; DeepSee, Spectra-Physics, Inc.) with a 25× water objective lens (XLPLN25XWMP2, NA 1.05). All imaging was performed by moving the motorized stages of the microscope along a virtual axis parallel to the optical axis of the objective lens. We acquired high-resolution anatomical stacks from anesthetized mice. Typical stacks consisted of 200-400 optical sections spaced 1 μm apart. The imaged area spanned 254 × 254 μm (512 × 512 pixels). To reduce the effects of the shot noise, a median filter (Fiji) was applied to the entire stack.

Analysis of mRNAs

The brain hemispheres were dissected from the mice and immediately frozen in liquid nitrogen. The tissue was homogenized, and total RNA was isolated using RNeasy (Qiagen) and RNase-Free DNase Set (Qiagen) according to the supplier's recommendations. To obtain cDNA, 2.5 μg of the total RNA was reverse transcribed with Primescript reverse transcriptase (Takara) using random primers in a 40-μl reaction volume.

qRT-PCR was performed using SYBR Premix ExTaq II (Takara) on a LightCycler480 (Roche). Primer sequences used for qRT-PCR are listed in Supplementary Table S1. All data were normalized to β2-microglobulin levels. The specificity of all primer pairs was confirmed by analyzing the melting curves.

Statistical analysis

Statistical analysis was performed using GraphPad Prism (version 6.0; GraphPad Software, La Jolla, CA, USA) and was performed using one-way ANOVA followed by Bonferroni's post hoc test or unpaired two-tailed Student's t-test, if applicable. All data are expressed as the mean ± S.E.M. Sample sizes were displayed in the figure legends. Statistical significance was defined as $p < 0.05$ and annotated as *$p < 0.05$, **$p < 0.01$, and ***$p < 0.001$. No significant difference is denoted as n.s.

**Supplementary Figures:**

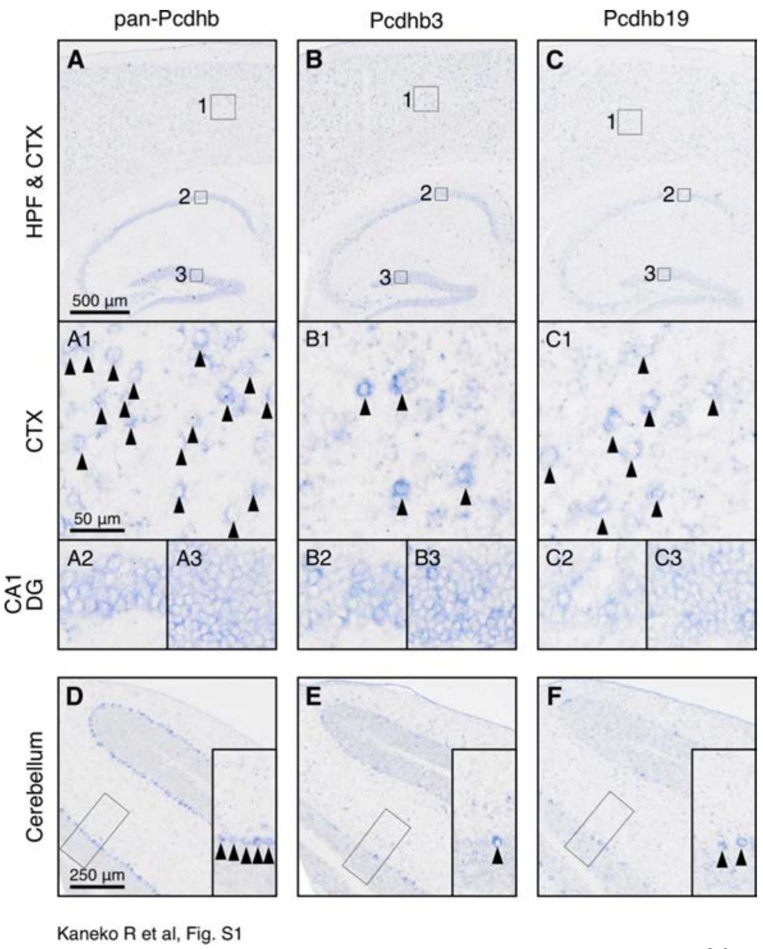


**Fig. S1. Distribution of endogenous cPcdhβ mRNAs in brain**

In situ hybridization for pan-cPcdhβ, cPcdhβ3 and cPcdhβ19 in wild-type mice at P28 is shown. The signals of pan-cPcdhβ transcripts were detected in the cerebral cortex and hippocampal CA1 region (A), or cerebellar Purkinje cells of the 6th cerebellar lobules (D). The scattered signals of cPcdhβ3 or cPcdhβ19 transcripts were detected in the cerebral cortex and hippocampal CA1 region (B-C), or cerebellar Purkinje cells of the 6th cerebellar lobules (E-F).

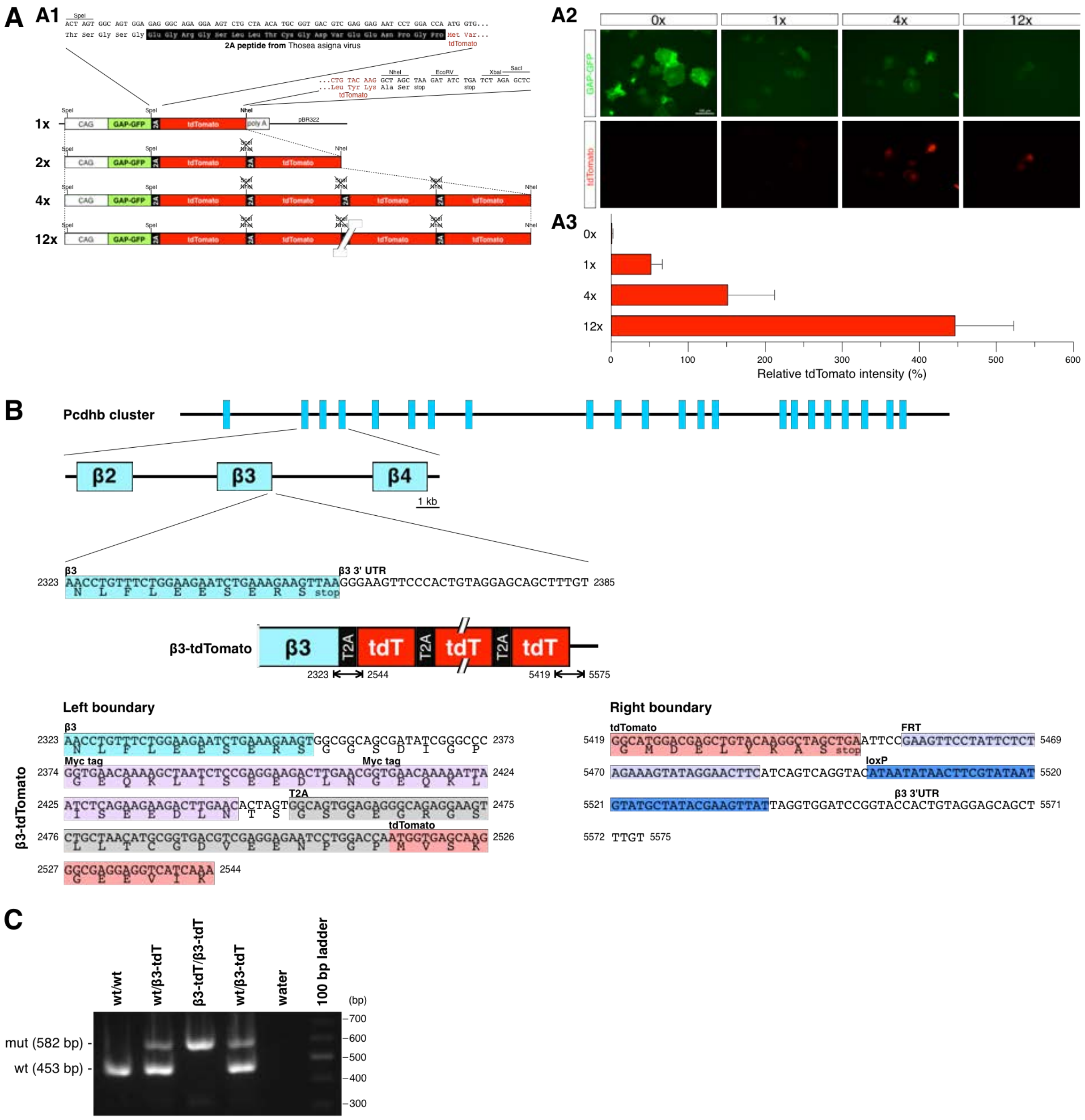


Kaneko et al, Fig S2

**Fig. S2. Generation of multiple T2A-tdTomato unit and cPcdhβ3 reporter mouse line**

(A) Enhancement of tdTomato fluorescence by multiple T2A-tdTomato. (A1) Design of multiple T2A-tdTomato plasmids. (A2) Representative images of Cos7 cells transfected with T2A-tdTomato plasmids. (A3) Relative tdTomato fluorescence intensity. The tdTomato fluorescence intensity was normalized to the GFP fluorescence intensity.

(B) Schematic representation of mouse cPcdhβ and cPcdhβ reporter loci. Genomic DNA sequences close to the β3-tdTomato knock-in site are shown.

(C) Genotyping PCR of β3-tdTomato mouse.

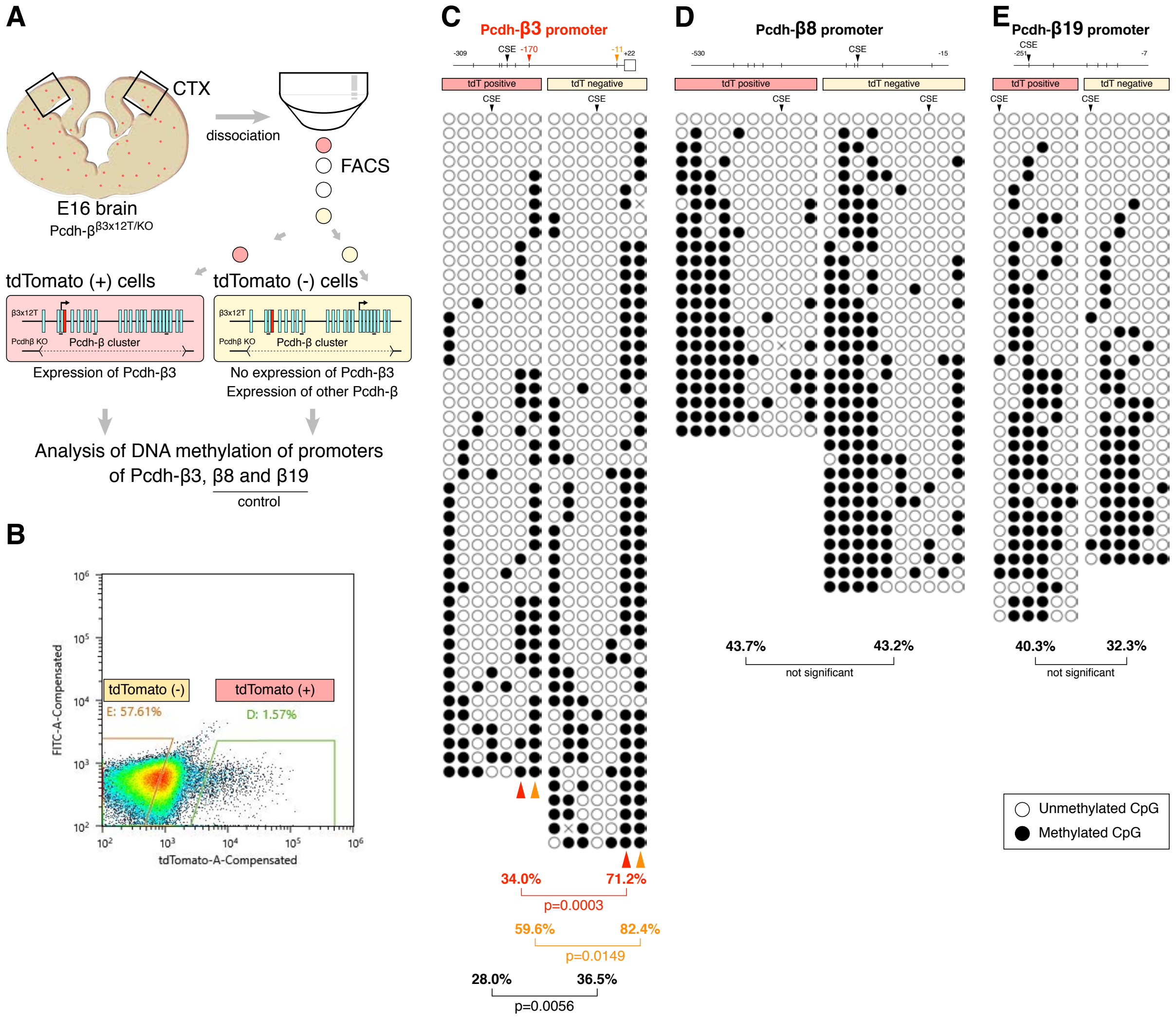


Kaneko R et al, Figure S3

**Fig. S3. Validation of β3-tdTomato reporter mouse by DNA methylation analysis of FACS-sorted tdTomato+ cells**

(A) Schematic drawing of the method. (B) Representative result for FACS-sorting of tdTomato+ cells. (C) Bisulfite sequencing of the cPcdhβ promoter regions in the tdTomato+ and tdTomato- cells. Each row represents a single DNA strand; the filled and open circles represent methylated and unmethylated CpG nucleotides, respectively. Average CpG methylation levels are indicated as a percentage.

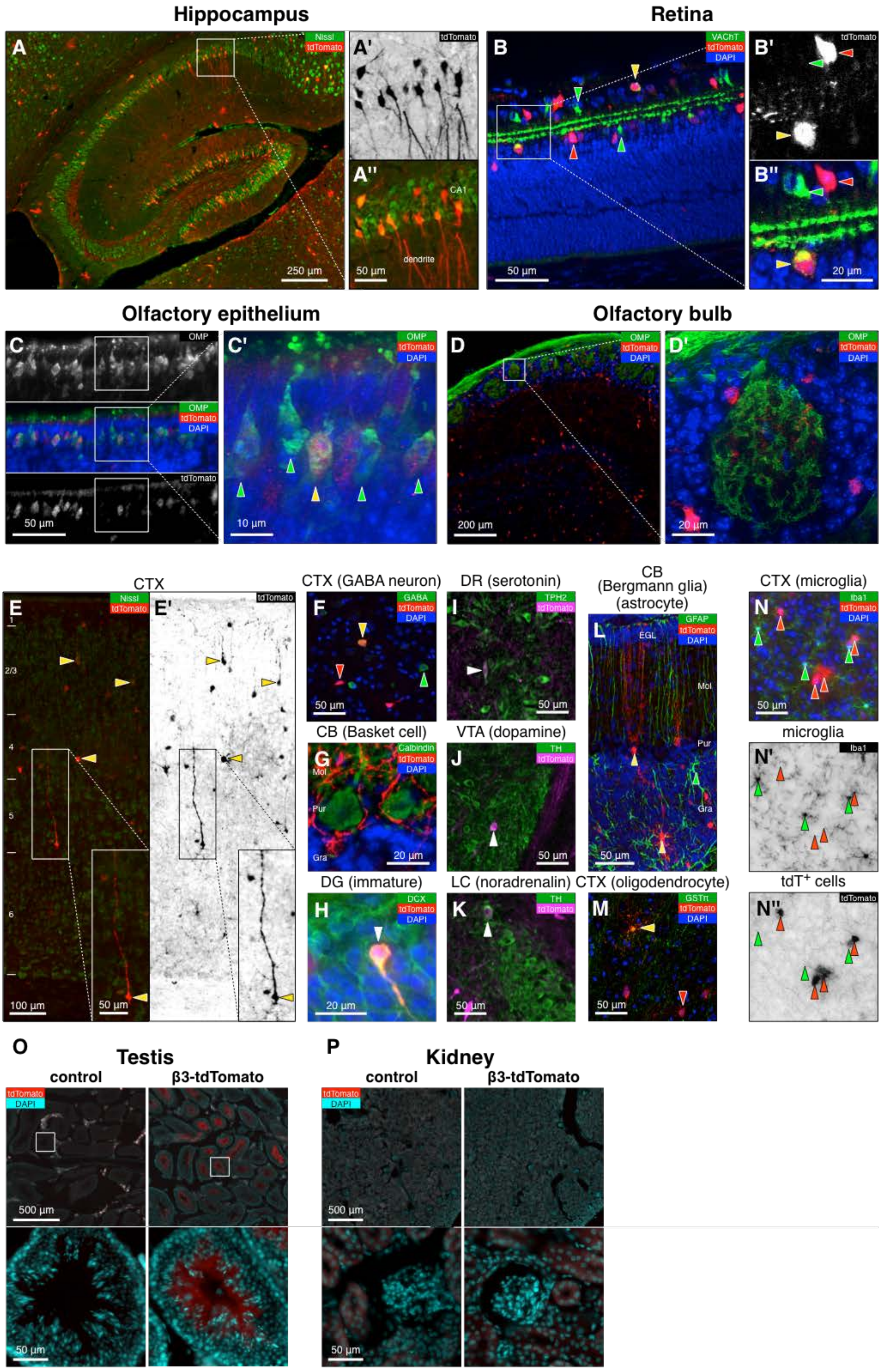


Kaneko R et al, Figure S3

**Fig. S4. Distribution of β3-tdTomato reporters in nervous systems are stochastic, but not in testis, and not detected in microglia and kidney**

(A-N) Representative image of β3-tdTomato reporter mouse nervous systems. (A) hippocampus at P9. (B) retina at P5. (C) olfactory epithelium at P0. (D) olfactory bulb at 4-week-old. (E) cerebral cortex at P7. (F) GABAergic neurons in cerebral cortex at 4w. (G) molecular layer interneuron in cerebellum at 4w. (H) immature neurons in hippocampal dentate gyrus at 4w. (I) serotonergic neurons in dorsal raphe nucleus at P7. (J) doapminergic neurons in ventral tegmental area at P7. (K) noradrenergic neurons in locus coellesence nucleus at P7. (L) Bergmann glia and astrocyte in cerebellum at 4w. (K) oligodendrocyte in cerebral cortex at 4w. (M) microglia in cerebral cortex at 4w.

(O)Representative images of adult testes from the control (left) and β3-tdTomato reporter mice (right).

(P) Representative images of adult kidneys from the control (left) and β3-tdTomato reporter mice mice (right).

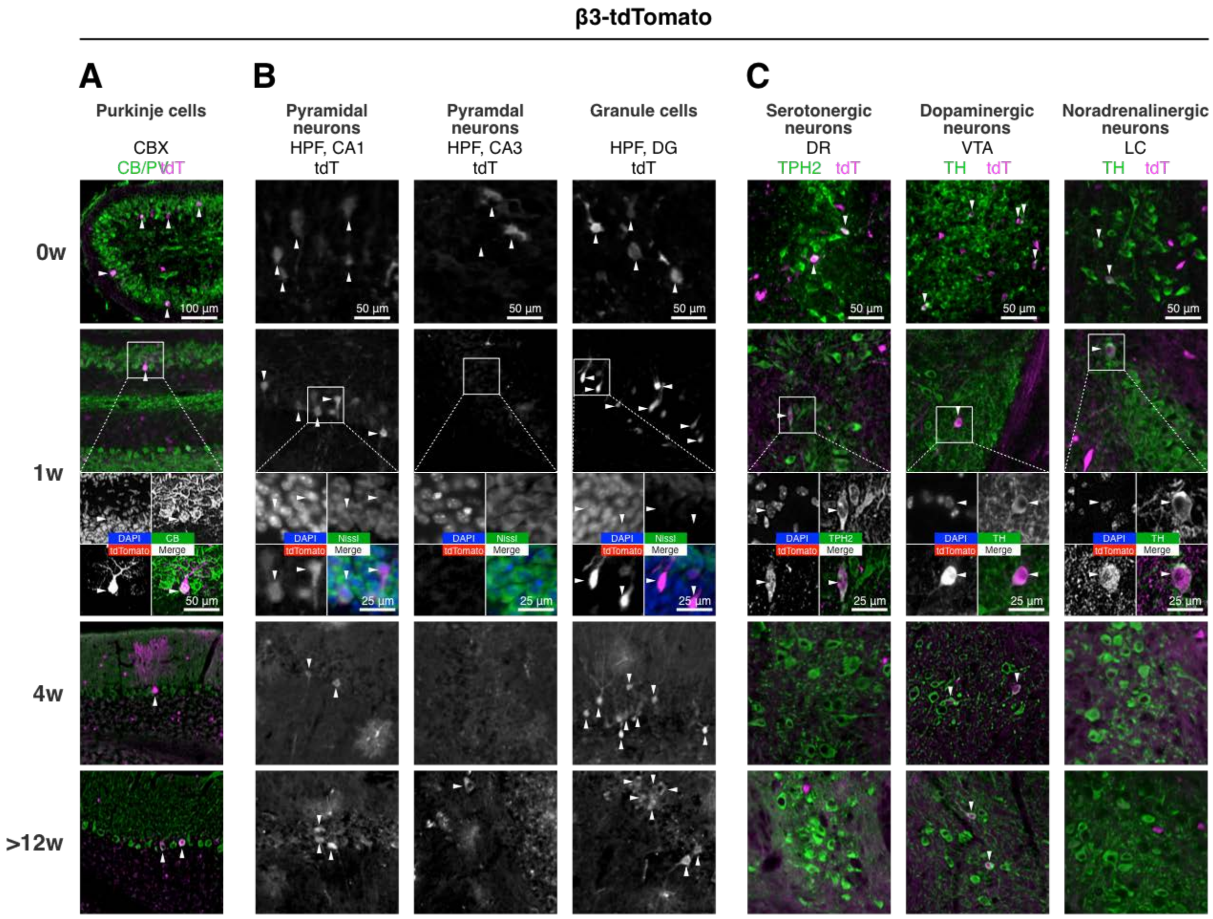


Kaneko R et al, Fig. S3

**Fig. S5. Developmental downregulation of stochastic cPcdhβ3 expression**

Representative images showing developmental downregulation of cPcdhβ3 expression frequency during postnatal maturation in the cerebellar Purkinje cells (A), hippocampal CA1 (B, left), CA3 (B, middle), and dentate gyrus (B, right), serotonergic neurons (C, left), dopaminergic neurons (C, middle), and noradrenalinergic neurons (C, right).

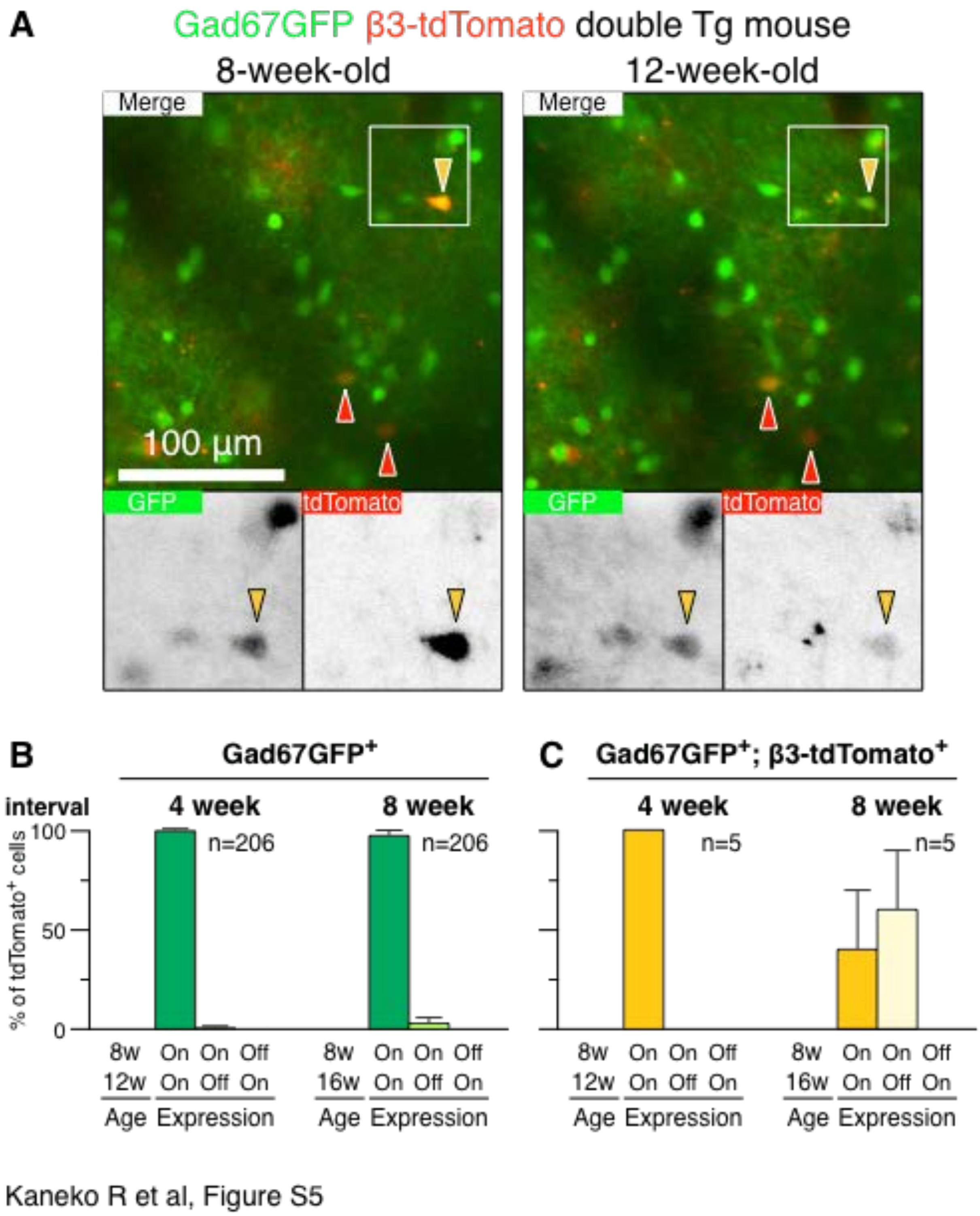


**Fig. S6. cPcdhβ3 expression in a single inhibitory neuron is dynamic**

The expression dynamics of cPcdhβ3 in living cerebral inhibitory neurons were assessed using in vivo two-photon microscopy. (A) Representative images of the somatosensory cortex of double-Tg mice harboring β3-tdTomato reporter allele and the Gad67-GFP mouse allele at 8-week-old (left) and the same region at 12-week-old (right). Lower panels show higher-magnification images of the boxed area showing the downregulation of β3-tdTomato in living inhibitory neurons (yellow arrowhead). (B) Quantification of the number of inhibitory neurons with unchanged GFP expression at both ages (green), switched expression from ON to OFF (light green), and from OFF to ON (white). Data obtained at 4-week intervals are shown in the left panel and at 8-week intervals in the right panel. (C) Quantification of the number of inhibitory neurons with unchanged β3-tdTomato expression at both ages (yellow), switched expression from ON to OFF (light yellow), and from OFF to ON (white). Data obtained at 4-week intervals are shown in the left panel and at 8-week intervals in the right panel. * $p<0.05$, Pearson’s chi-square test.

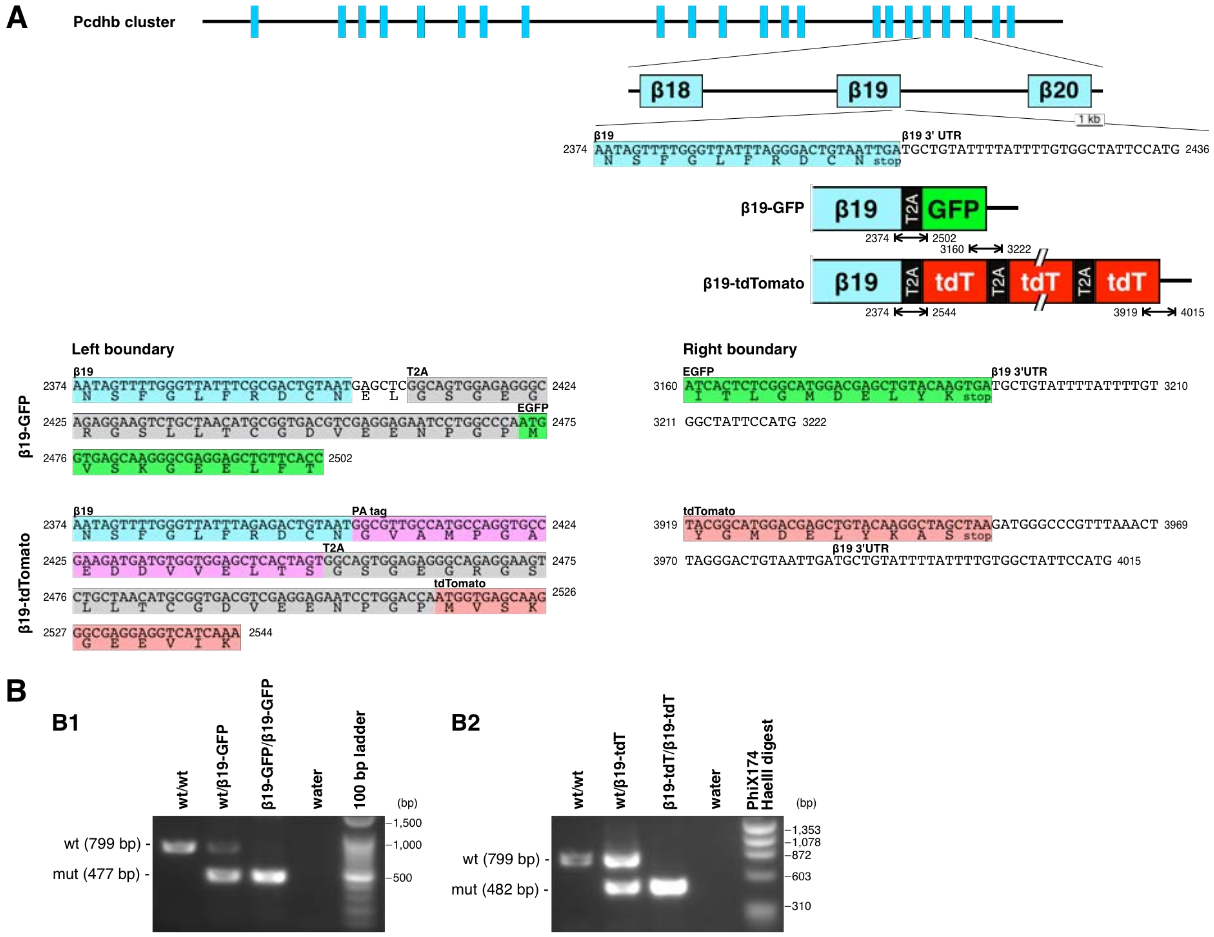


Kaneko et al, Figure S6

**Fig. S7. Generation of cPcdhβ19-GFP and cPcdhβ19-tdTomato reporter mouse lines**

(A)Schematic representation of mouse cPcdhβ and cPcdhβ reporter loci. Genomic DNA sequences close to the β19-GFP and β19-tdTomato knock-in site are shown.

(B)Genotyping PCR of β19-GFP mouse (B1) and β19-tdTomato mouse. (B2)

### β19-tdTomato

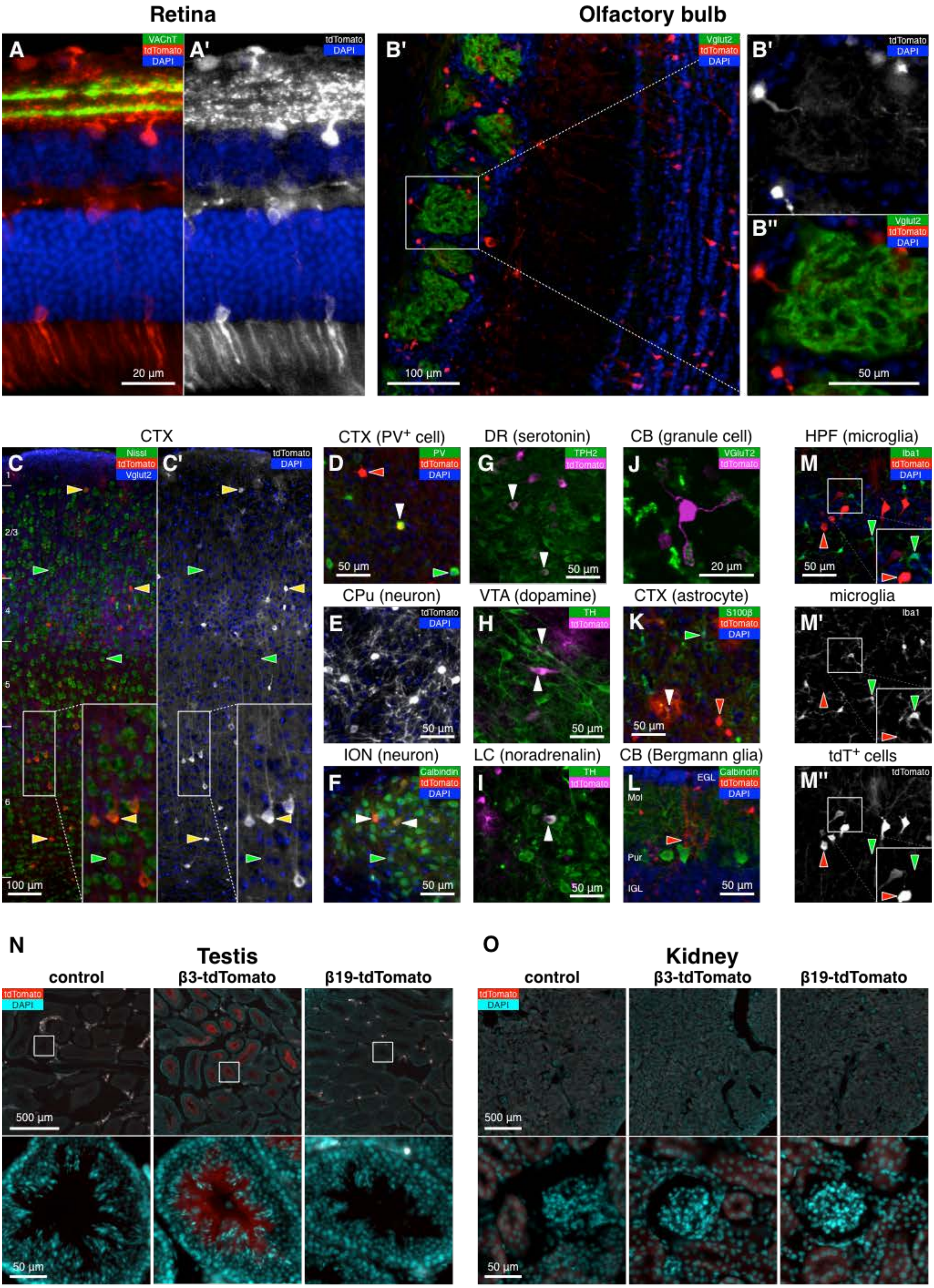


Kaneko R et al, Figure S7

**Fig. S8. Distribution of β19-tdTomato reporters in nervous systems are stochastic, but not detected in microglia, testis, and kidney**

(A-M) Representative image of β19-tdTomato reporter mouse nervous systems. (A) retina at P28. (D) olfactory bulb at 4-week-old. (C) cerebral cortex at P7. (D) Parvalbumin$^+$ neurons in cerebral cortex at 4w. (E) neurons in caudate putamen at 4w. (F) neurons in inferior olivary nucleus s at 4w. (G) serotonergic neurons in dorsal raphe nucleus at P7. (H) doapminergic neurons in ventral tegmental area at P7. (I) noradrenergic neurons in locus coellesence nucleus at P7. (J) granule cell in cerebellum at 4w. (K) astrocyte in cerebral cortex at 4w. (L) Bergman glia in cerebellum at 4w. (M) microglia in hippocampus at 4w.

(N)Representative images of adult testes from the control (left), β3-tdTomato reporter mice (middle), and β3-tdTomato reporter mice (right). The images of control and β3-tdTomato reporter mice are identical with Fig. S3.

(O) Representative images of adult kidneys from the control (left), β3-tdTomato reporter mice (middle), and β3-tdTomato reporter mice (right). The images of control and β3-tdTomato reporter mice are identical with Fig. S3.

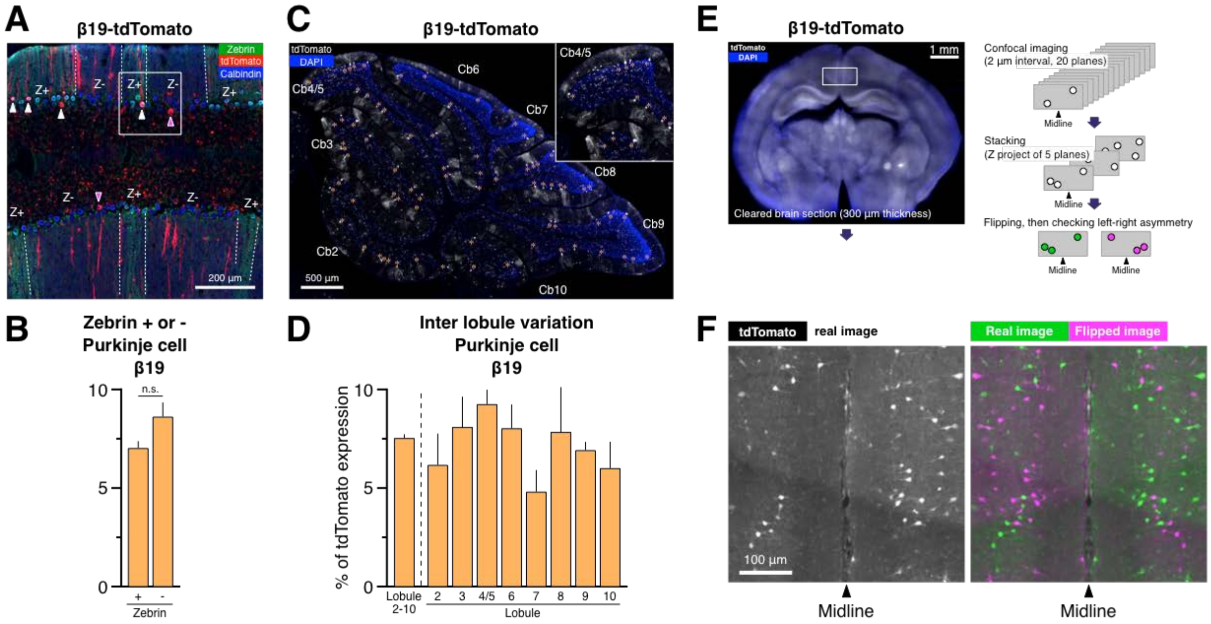


Kaneko R et al, Fig. S6

**Fig. S9. Spatial distribution β19-tdTomato reporter in the cerebellum and frontal cortex is stochastic**

(A-C) Representative cerebellar section of β19-tdTomato reporter mice. Coronal section of β19-tdTomato reporter mouse cerebellum (A)(counterstained with DAPI), with the boxed region magnified in (B)(identical with Fig. 2E) showing stochastic tdTomato expression in Zebrin II-positive (labeled with Zebrin II and calbindin) and Zebrin II-negative Purkinje cells (labeled with calbindin)(C).

(D-F) Left-right asymmetry of cPcdhβ19 expression in the frontal cortex. (D) Schematic drawing of the method used to analyze left-right asymmetry of cPcdhβ19 expression. In brief, an imaging method to clear 300-µm-thick brain sections with SeeDB2, to image at 20x on the DragonFly confocal microscopy, to digitally reconstruct, and to examine left-right asymmetry of the distribution of tdTomato$^{+}$ cells in the nearby midline of the cerebral cortex. (E) Representative z-stack images of β19-tdTomato reporter nearby the midline of the frontal cortex. (F) Random and left-right asymmetric distribution of cPcdhβ19 expression. Left panel shows real images of the regions indicated in (E). Right panel shows merged images of the real image (green) and horizontally-flipped image (magenta).

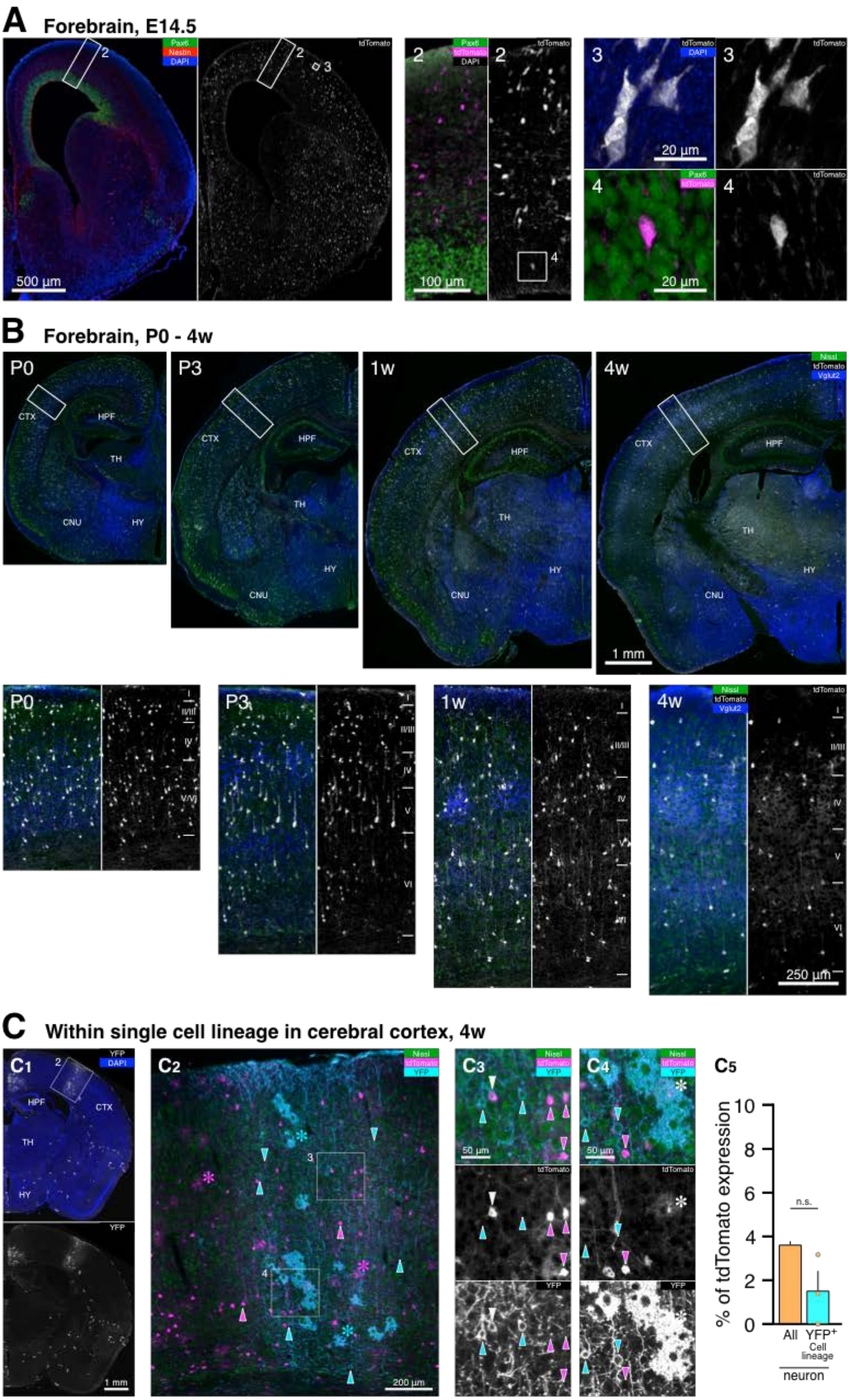


Kaneko R et al, Fig. S7

**Fig. S10. cPcdhβ19 expression in developing neurons is dynamic at the population level**

(A) Representative images showing stochastic cPcdhβ19 expression in the embryonic cortex.

(B) Representative images showing developmental downregulation of cPcdhβ19 expression frequency in the somatosensory cortex during postnatal maturation.

(C) Stochastic cPcdhβ19 expression within single cell lineage in the cerebral cortex. (B1-B4) Representative image of triple Tg mouse harboring β19-dTomato, TFC.09, and Cre-dependent channelrhodopsin reporter (Ai32), showing stochastic cPcdhβ19 expression within a single cell lineage in the cerebral cortex. The higher-magnification images of the boxed area in B1 and B2 are shown. (B5) Frequency of tdTomato$^{+}$ cells among total neuron or YFP-positive neurons.

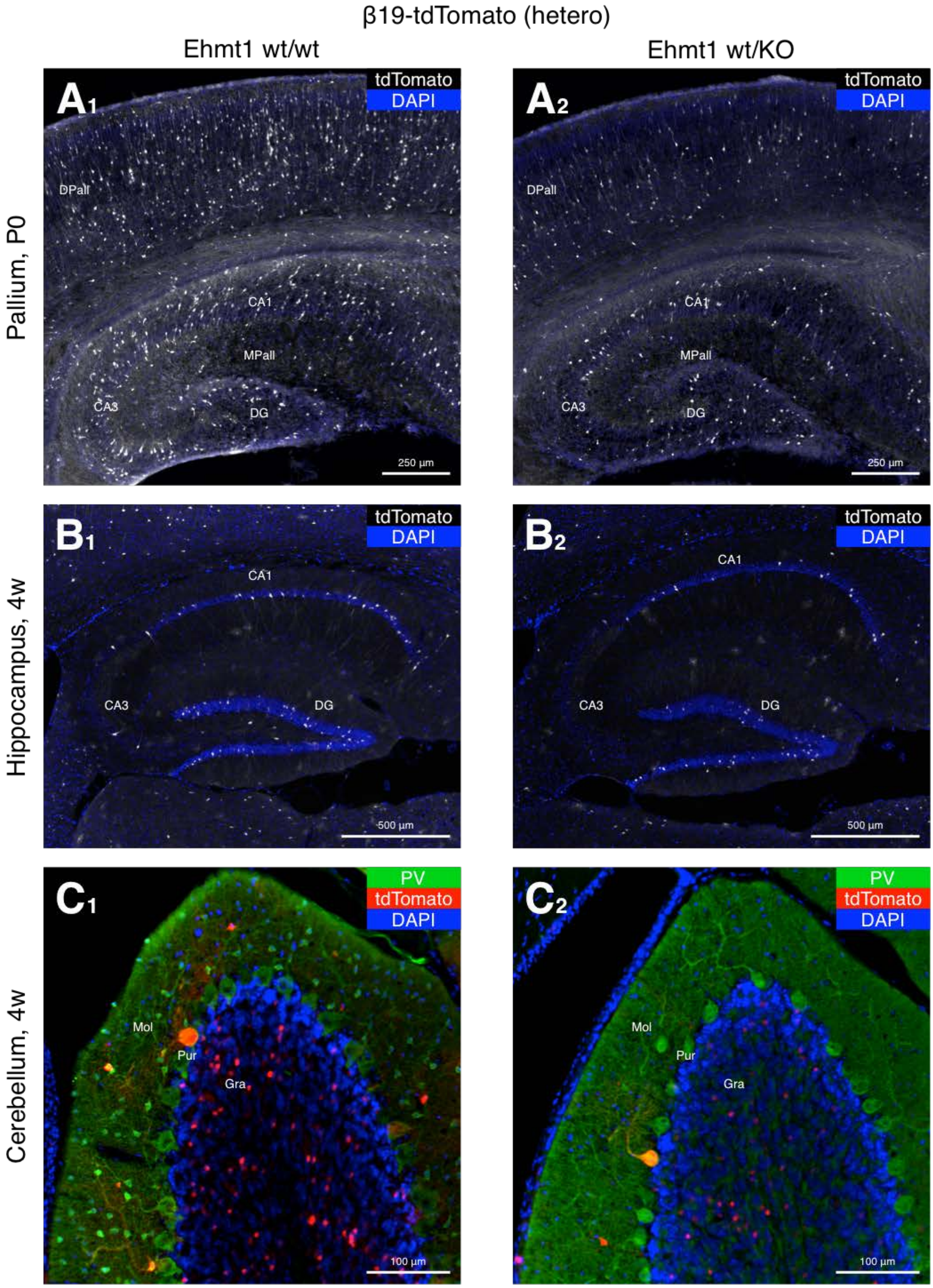


Kaneko R et al, Fig. S8

**Fig. S11. Dysregulation of cPcdhβ19 expression in intellectual disability model**

Expression of cPcdhβ3 and cPcdhβ19 is downregulated in a mouse model of intellectual disability (KS). (A) Representative images of the sagittal section of the whole brain (upper panels) and the dorsal and medial pallium (isocortex and hippocampal allocortex)(lower panels) in control and Ehmt1$^{+/-}$ β3-tdTomato reporter mice on postnatal day 0. (B) Representative images of sagittal sections of the dorsal and medial pallium (isocortex and hippocampal allocortex) from control and Ehmt1$^{+/-}$ β19-tdTomato reporter mice on postnatal day 0.
